# A release-and-capture mechanism generates an essential non-centrosomal microtubule array during tube budding

**DOI:** 10.1101/2020.05.21.108027

**Authors:** Ghislain Gillard, Gemma Girdler, Katja Röper

**Affiliations:** MRC-Laboratory of Molecular Biology, Francis Crick Avenue, Cambridge Biomedical Campus, Cambridge CB2 0QH, UK; Wellcome-MRC Stem Cell Institute, Jeffrey Cheah Biomedical Centre, Cambridge Biomedical Campus, University of Cambridge, Puddicombe Way, Cambridge, CB2 0AW, UK

## Abstract

Non-centrosomal microtubule arrays serve crucial functions in cells, yet the mechanisms of their generation are poorly understood. During budding of the epithelial tubes of the salivary glands in the *Drosophila* embryo, we previously demonstrated that the activity of pulsatile apical-medial actomyosin depends on a longitudinal non-centrosomal microtubule array. Here we uncover that the exit from the last embryonic division cycle of the epidermal cells of the salivary gland placode leads to the mother centrosome in the cells losing all microtubule-nucleation capacity. This restriction of nucleation activity to the daughter centrosome is key for proper morphogenesis. Furthermore, the microtubule-severing protein Katanin and the minus-end-binding protein Patronin accumulate in an apical-medial position only in placodal cells. Loss of either in the placode prevents formation of the longitudinal microtubule array and leads to loss of apical-medial actomyosin and apical constriction. We thus propose a mechanism whereby Katanin-severing at the single active centrosome releases microtubule minus-ends that are then anchored by apical-medial Patronin to promote formation of the longitudinal microtubule array crucial for apical constriction and tube formation.

## Introduction

The microtubule cytoskeleton plays many essential roles in cells, from faithful chromosome segregation during cell division to transport of many cargoes. In most animal cells that are actively dividing and cycling, the microtubule cytoskeleton is nucleated and anchored at centrosomes throughout interphase but especially during mitosis ^1^. Centrosomes consist of a single or a pair of centrioles at their core, depending on the stage of the cell cycle, surrounded by a cloud of pericentriolar material (PCM) that contains the critical microtubule nucleator γ-tubulin in form of the γ-tubulin ring complex (γ-TURC) that templates the microtubule protofilament arrangement (Fig.2A; ^2^).

However, in post-mitotic cells such as neurons and epithelial cells, microtubules can also be nucleated from non-centrosomal sites. Non-centrosomal microtubule function in those cells is crucial for processes such as directed intracellular transport, organelle positioning and cell polarity ^3-5^. In post-mitotic epithelial cells non-centrosomal microtubule can be organised in different arrays, lying for instance parallel to the apical surface ^6, 7^ or forming extended longitudinal arrays along the apical-basal axis ^8, 9^. There is now growing evidence for a role of microtubules in epithelial morphogenesis. They can do so by exerting forces against the plasma membrane ^10^, by coordinating forces at the tissue-scale level ^6, 7^ or by regulating acto-myosin localisation or activity ^8, 11^. Despite such important cellular and developmental functions of non-centrosomal microtubules it remains unclear, though, what the mechanism of non-centrosomal microtubule generation is, whether it involves for instance nucleation from non-centrosomal MTOCs, or whether pre-existing microtubules become relocalised ^12^.

We have previously shown a function for a longitudinal non-centrosomal microtubule array during tube morphogenesis in the *Drosophila* embryo ^8^. During the initial tissue bending and budding process of the tubes of the salivary gland from an epithelial placode (Fig. 1A-A’’), microtubules rearrange by 90° from a previously apical centrosomal to a longitudinal non-centrosomal array. This happens concomitantly with the cells undergoing apical constriction (Fig.1A’’), which itself is required to drive the budding morphogenesis ^8, 13^. Disruption of the microtubule cytoskeleton leads to the selective loss of an apical-medial, but not junctional, pool of actomyosin, and this pool is required for successful apical constriction ^8^. It is thus far unclear what triggers and establishes the microtubule rearrangement and generation of the non-centrosomal array.

**Figure 1.**
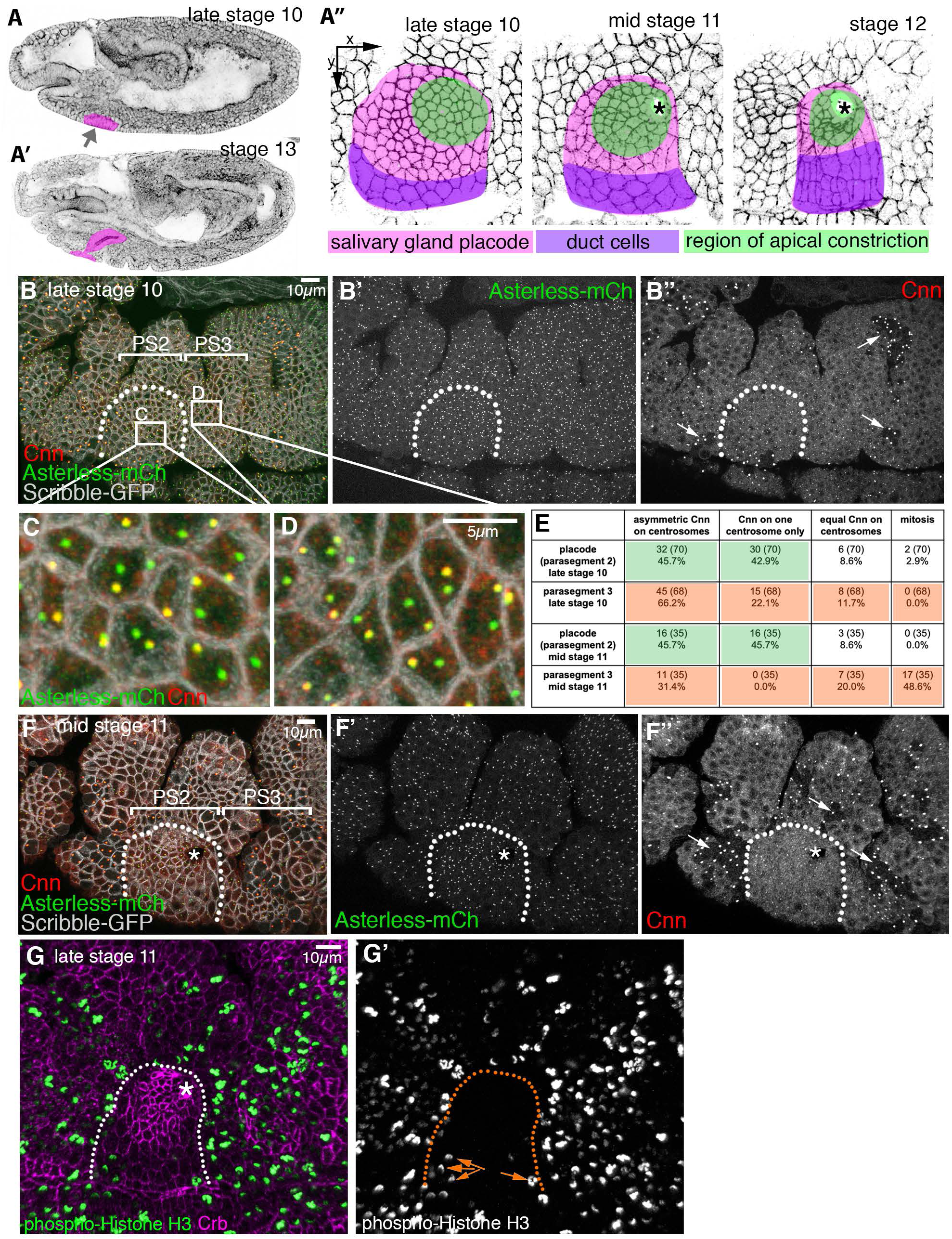
Changes to centrosomes during tube budding. **A-A’’** The salivary glands form from two epithelial placodes localised on the ventral surface of the *Drosophila* embryo that become specified at the end of embryonic stage 10 (**A**). These placodes invaginate through budding to form a simple tube (**A’**). **A’’** During invagination, cells close to the forming invagination point (asterisks) in the dorsal-posterior corner constrict apically (green cells) as part of the morphogenetic programme ^13^. Pink marks all secretory cells of the placode, magenta marks future duct cells near the ventral midline. **B-F’’** Salivary gland placodal cells in embryonic parasegment 2 (**B, F**) are the first epithelial tissue primordium to enter G_1_ phase of embryonic division cycle 17. This post-mitotic state leads to a permanent loss of Cnn (**B’’, F’’**) from one centrosome after the last mitosis (quantification in **E**). Arrows in **B’’** and **F’’** point to mitotic domains in the epidermal tissue surrounding the placode. **C** and **D** illustrate the permanent (PS2) and temporary (PS3) Cnn asymmetry in cells; note that epidermal cells in the domain shown in **D** have entered mitosis again in **F’’**. **G-G’** Phospho-histone H3 labeling (green and single channel in **G’**) to mark mitotic cells at late stage 11 clearly shows that the secretory cells of the salivary gland placode have stopped dividing, only a few duct cell precursors near the ventral midline still undergo mitoses (orange arrows in **G’**), whereas epidermal cells in the surrounding epidermis are still undergoing division cycle 16. The salivary gland placode is indicated by dotted lines and the invagination point, where present, by an asterisk. See also Figure S1.

*Drosophila* embryos undergo a modified fast cell cycle in early embryogenesis ^14^. At the beginning of tissue morphogenesis, divisions become asynchronous and most cells in the embryo only divide three more times, ^15^. This leads the epidermal cells after each M-phase in these cycles to inherit two separated centrosomes consisting of a mother centriole with an attached pro-centriole or daughter ^16^. At the time point that salivary gland morphogenesis commences in the embryo, the salivary gland placodal cells differ from all other epidermal cells in that they are the first to finish all embryonic division cycles and are the first cells to enter a G1 phase in interphase of embryonic cell cycle 17 and concomitantly become post-mitotic ^15^. At the end of the last mitosis 16, in contrast to the previous cycles, these cells now inherit two centrosomes consisting of an isolated centriole each, with no pro-centriole or daughter ^16^.

Here, we describe our discovery of a step-wise process that implements these changes in the salivary gland placode: as part of concluding embryonic mitoses, the cells of the placode are the first to enter a G_1_ phase with concomitant loss of microtubule nucleation capacity of the mother centrosome, a loss that we show is important to ensure proper morphogenesis. Furthermore, microtubules generated from the remaining active daughter centrosome are released by severing through Katanin and then anchored and stabilised by Patronin, the *Drosophila* CAMSAP homologue ^9, 17, 18^. Both Katanin and Patronin function, we show, are required for microtubule rearrangements, for the proper apical-medial actomyosin activity and apical constriction during tube budding morphogenesis.

## Results

### Changes to centrosomal microtubule-nucleation capacity in the placode

In order to investigate whether changes at centrosomes contributed to the formation of the non-centrosomal microtubule array in the salivary gland placode, we decided to analyse levels of centrosome components. The overall capacity and requirement for centrosomal versus non-centrosomal microtubule nucleation during *Drosophila* embryogenesis, beyond the fast synchronised cell cycles 1-13 at very early stages, is unclear. Maternal loss of key centrosome components leads to developmental arrest of early embryos during the fast syncytial divisions ^19^, whereas in zygotic mutants the protein levels of centrosome components only run out in larval stages and embryogenesis is unaffected ^20^. Upon specification, the cells of the salivary gland primordium, the placode (Fig. 1A, A’’), have completed mitosis 16 and cease full mitoses for the remainder of embryogenesis. They enter their first G_1_ phase during nuclear cycle 17 and remain in it until the end of embryogenesis, hence being post-mitotic ^16^. Amongst cells of the anterior ectoderm, the salivary gland placodal cells are in fact the first organ primordium to reach this stage (Fig. 1B-B’’, F-F’’, G-G’). Concomitantly, salivary gland placodal cells now enter endoreduplication or endocycles, i.e. DNA-synthesis and segregation without any accompanying cytokinesis, leading to most salivary gland cells being polytene at larval stages ^21^.

Interestingly, labelling of centrosomes within the salivary gland placode at late stage 10, just prior to start of the budding morphogenesis, revealed that most post-mitotic placodal cells contained two well-separated centrosomes that showed a striking asymmetry in the accumulation of Centrosomin (Cnn), a component of the PCM key to centrosome maturation (Fig.1B-E; 45.7% of cells with an asymmetry of Cnn accumulation and another 42.9% with Cnn completely restricted to one centrosome) ^22, 23^. These placodal cells, that will form the secretory part of the gland (Fig. 1A’’), are in G_1_ phase of cycle 17 ^16^. Most cells in the surrounding epidermis at this stage have not completed cell divisions, reflected by the continuing occurrence of mitoses and accompanying clusters of high mitotic Cnn labeling (Fig. 1B’’, F’’ arrows), as the microtubule nucleation capacity of centrosomes increases dramatically during mitosis, concomitant with a drastic increase in PCM ^24^. Actively dividing cells just after M-phase will also show Cnn asymmetry as the daughter centrosome upon division needs to re-recruit PCM prior to the next S-phase ^25^. Hence, many cycling epidermal cells also displayed a temporary Cnn asymmetry (Fig. 1D, E; analysed in parasegment 3 (PS3): 66.2% of cells with asymmetric Cnn accumulation and 22.1% of cells with Cnn restricted to one centrosome). Such epidermal asymmetry was evident from stage 9 onwards (Fig.S1), representing cells in G2 of cycle 15 or 16 that are in the process of PCM recruitment about to generate two centrosomes with equal Cnn prior to the next M-phase. Accordingly, at mid stage 11, half of the epidermal cells in parasegment 3 had entered mitosis again (48.6%) and 20.0% in addition showed equal Cnn distribution (Fig. 1E-F’’), thus most likely being in G_2_ just prior to the next mitosis. In contrast, in the already post-mitotic placodal cells the Cnn asymmetry remained identical (Fig. 1E). Note that cells close to the ventral midline, that will later form part of the duct of the salivary gland (Fig. 1A’’), re-entered mitosis once more, as evident by phospho-histone H3 labeling at late stage 11 (Fig. 1G, G’). In contrast to Cnn, Asterless (Asl), a core component of the centriole, was equally enriched on both centrosomes throughout the epidermis and placode (Fig. 1B-F’).

We also investigated the distribution of other centrosome components in placodal cells and found that the core centriole components Asl, Spd-2 and Sas-4 (Fig. 2A) were equally enriched on both centrosomes (Fig. 2 B-C’). However, in addition to the asymmetric distribution of Cnn, γ-tubulin, that is recruited by Cnn ^26^, and Polo-kinase, a key regulator of PCM recruitment to the centrosome ^22, 27^, were enriched asymmetrically (Fig. 2B-C’ and Fig. S2). This asymmetry explained the lack of nucleation capacity of some of the centrosomes analysed in live assays using EB1-GFP to label growing microtubule plus-ends and Asl-mCherry (Fig. S2A-C). Consistent with the asymmetry in nucleation capacity, 90% of centrosomes with higher levels of Cnn within the placode actively nucleated microtubules in live assays (Fig. 2D-E), underlining that only Cnn-enriched centrosomes still retain nucleation capacity. Therefore, the post-mitotic placodal cells displayed a pronounced and permanent centrosome asymmetry and hence asymmetry in the capacity to nucleate microtubules.

**Figure 2.**
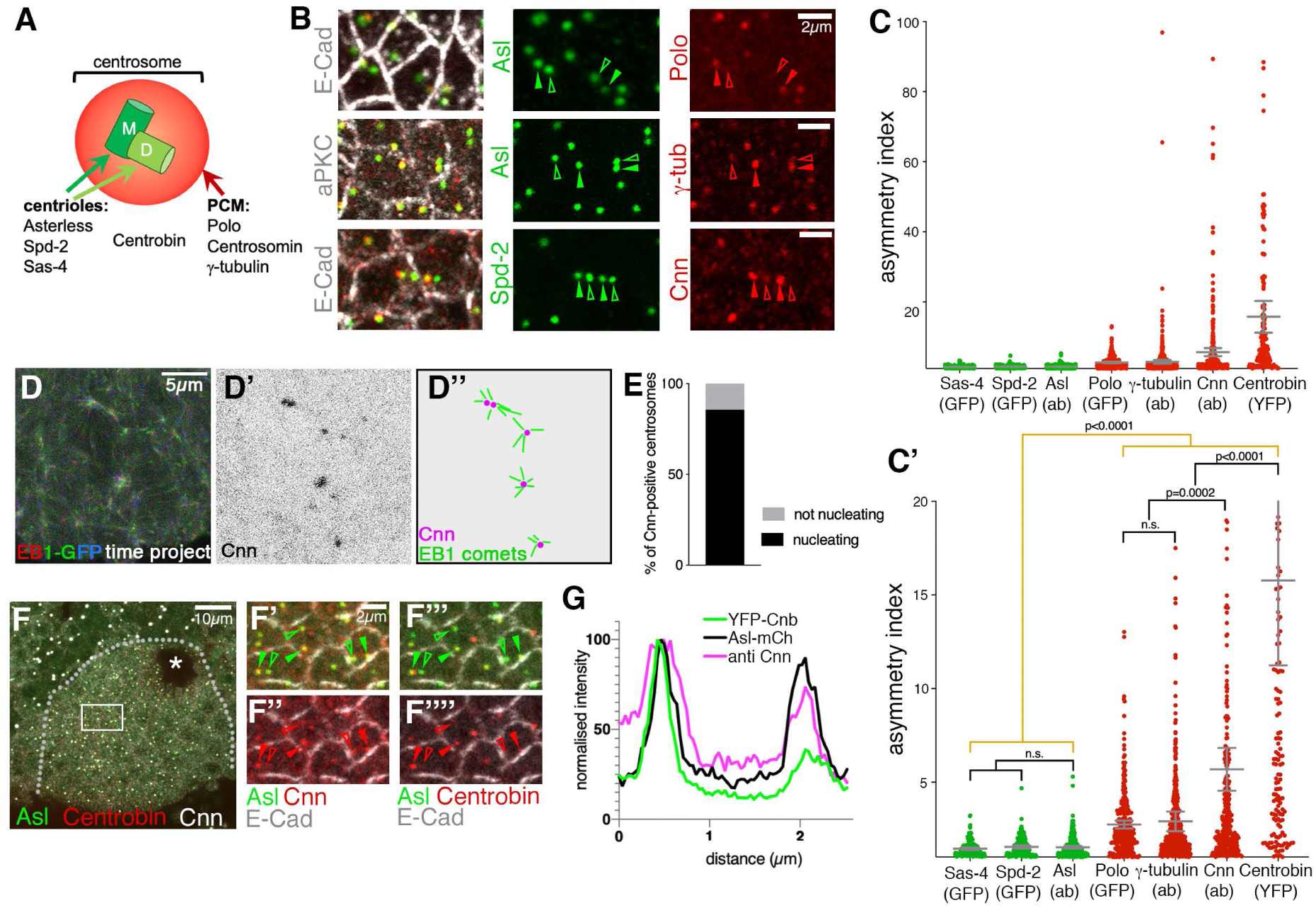
Placodal centrosomes show strong asymmetry of PCM constituents and microtubule nucleation capacity. **A** Centrosomes are usually built of two centrioles, a mother centriole (M) inherited from the last division, and a newly nucleated daughter centriole (D), together surrounded by a cloud of PCM. Key centriole components include Asl, Spd-2 and Sas-4, whereas the PCM contains components such as Polo-kinase, Cnn and γ-tubulin. The centriole component Centrobin is only found on the daughter centriole. **B-C’** Whereas the centriole components Asl, Spd-2 and Sas-4 are equally enriched on both centrosomes in placodal cells (localisation for Asl and Spd-2 in **B**, quantification for all in **C, C’**), PCM components Polo, Cnn and γ-tubulin show asymmetric accumulation at placodal centrosomes (localisation in **B** and quantification in **C, C’**). Cell outlines are marked by E-Cadherin (E-Cad) or aPKC in **B**. **C, C’** n values are: *Sas4-GFP*: 170 cells from 5 embryos; *Spd-2-GFP*: 139 cells from 4 embryos; Asl (antibody, ab): 3302 cells from 9 embryos; *GFP-Polo*: 270 cells from 7 embryos; γ-tubulin (ab): 487 cells from 12 embryos; Cnn (ab): 309 cells from 9 embryos; *Cnb-YFP*: 157 cells from 5 embryos. Pairwise comparison of distributions was done via Kruskal-Wallis test (one-way ANOVA), p values and significance are indicated in **C’**. Mean and 95% CI are shown. **D-E** 90% of Cnn-containing centrosomes actively nucleate microtubules in salivary gland placodal cells. **D** RGB-projection of 3 consecutive time frames, 1s apart, of a time lapse movie of *EB1-GFP Cnn-RFP* flies, showing the EB1-GFP channel only. **D’** shows the Cnn-RFP channel to indicate the positions of centrosomes. **D’’** schematically illustrates the EB1 ‘comets’ moving away from Cnn-positive centrosomes that indicate active microtubule nucleation. **E** Quantification of nucleation capacity of 70 Cnn-positive centrosomes. **F-G** Cnn and the daughter-centrosome marker Centrobin (Cnb) accumulate asymmetrically in individual cells, but on the same centrosome. Box in **F** is shown enlarged in **F’-F’’’’**. Asl labels all centrosomes and E-Cad labels cell outlines in **F’-F’’’’. G** Line scan profile through both centrosomes of a single cell to illustrate the co-enrichment of Cnb and Cnn on the same centrosomes. Note that Cnb asymmetry is much more pronounced than Cnn asymmetry, as also quantified in **C, C’**. More line scan examples are shown in Figure S2. The salivary gland placode is indicated by a dotted line and the invagination point by an asterisk in **F**. Filled arrowheads in **B** and **F’-F’’’’** indicate centrosomes with PCM and Centrobin accumulation, respectively, while hollow arrowheads point towards centrosomes without PCM or Centrobin staining.

How is this interphase asymmetry in centrosomes achieved? Similar to the salivary gland placode, in *Drosophila* larval neuroblast only one centrosome (composed at this stage of a single centriole as in the salivary gland placode) nucleates an interphase array, and this is the daughter centrosome/centriole ^28^. This centriole is selectively marked by the daughter-specific centriole component Centrobin ^29^. In neuroblasts, upon finishing a mitosis, the mother centriole/centrosome initially contains more PCM including Cnn, but this is then lost over time whilst the daughter centriole/centrosome that contains Centrobin continues to accumulate more and more PCM ^30^. To test whether a similar mechanism could be at work in the placode, we analysed the distribution of Centrobin, using *Ubi::Centrobin-YFP* ^29^, in the placodal cells (Fig. 2C, F-G). Centrobin was highly asymmetric (Fig. 2C,C’) and was always enriched on the one centrosome in placodal cells that also contained higher levels of Cnn, as visualised in line scans through centrosomes of a single cell (Fig. 2G, Fig. S2D-D’’’). Polo-kinase, γ-tubulin and Cnn were as expected enriched on the same centrosome in placodal cells, though the Polo and γ-tubulin asymmetry were not as pronounced as the difference in Cnn (Fig. 2C’, Fig. S2E-J).

Taken together these result show that, as in neuroblasts, only the daughter centrosome in the secretory cells of the salivary gland placode during early tube morphogenesis retains microtubule nucleation capacity.

### Centrosome asymmetry is a prerequisite for proper morphogenesis

We next sought to determine whether centrosome asymmetry and restriction to nucleation from a single centrosome was important for the generation of the non-centrosomal array or the morphogenesis. To address this question, we analysed embryos overexpressing a transgene of γ-tubulin expressed under the *ncd* promoter that is expressed at high levels up to mid-embryogenesis (*ncd::γ-tubulin-EGFP*; Fig. 3A-B). In such embryos the γ-tubulin centrosomal asymmetry in placodal cells was less pronounced (Fig. 3A’, B). The whole embryonic epidermis where *ncd* is expressed showed an increase in microtubule intensity compared to controls. This was particularly pronounced in the salivary gland placodal cells overexpressing γ-tubulin-EGFP, as measured through labelling of either α-tubulin (Fig. 3C and Fig. S3A-B’) or tyrosinated α-tubulin labeling (Fig. 3C-D’). Labelling for stable microtubules (using acetylated α-tubulin staining) did not increase but was rather slightly reduced (Fig.3C, E, E’), suggesting the overall increase reflected newly polymerised microtubules. As the secretory cells are undergoing apical constriction at this time point, we also analysed actomyosin levels in embryos. In addition to an increase in overall microtubule intensity, levels of F-actin also increased strongly, and this was again particularly enhanced in the placodal future secretory cells for both junctional and apical-medial actin (Fig. 3F, F’ and Fig. S3E). Similarly, myosin II levels (revealed by an endogenously tagged version of myosin regulatory light chain, Sqh-RFP; ^31^) were increased (Fig. 3M and Fig. S3C-D’). Furthermore, cell apices in *ncd::γ-tubulin-EGFP* placodes often showed a convoluted apical junctional morphology (‘wavy’ junctions; Fig. 3H’, I-K,) that is indicative of strong, possibly excessive, apical constriction ^32, 33^. In addition, cells showed more constricted apices compared to control (Fig. 3 G, H, L). Excessive amounts of microtubules generated in placodal cells of *ncd::γ-tubulin-EGFP* embryos, though nucleated at centrosomes, in the placodal cells appeared to contribute to an enlarged non-centrosomal array (Fig. 3 N, O), which we suggest is the underlying cause of the increase in apical-medial actomyosin and increased apical constriction (Fig. 3P).

**Figure 3.**
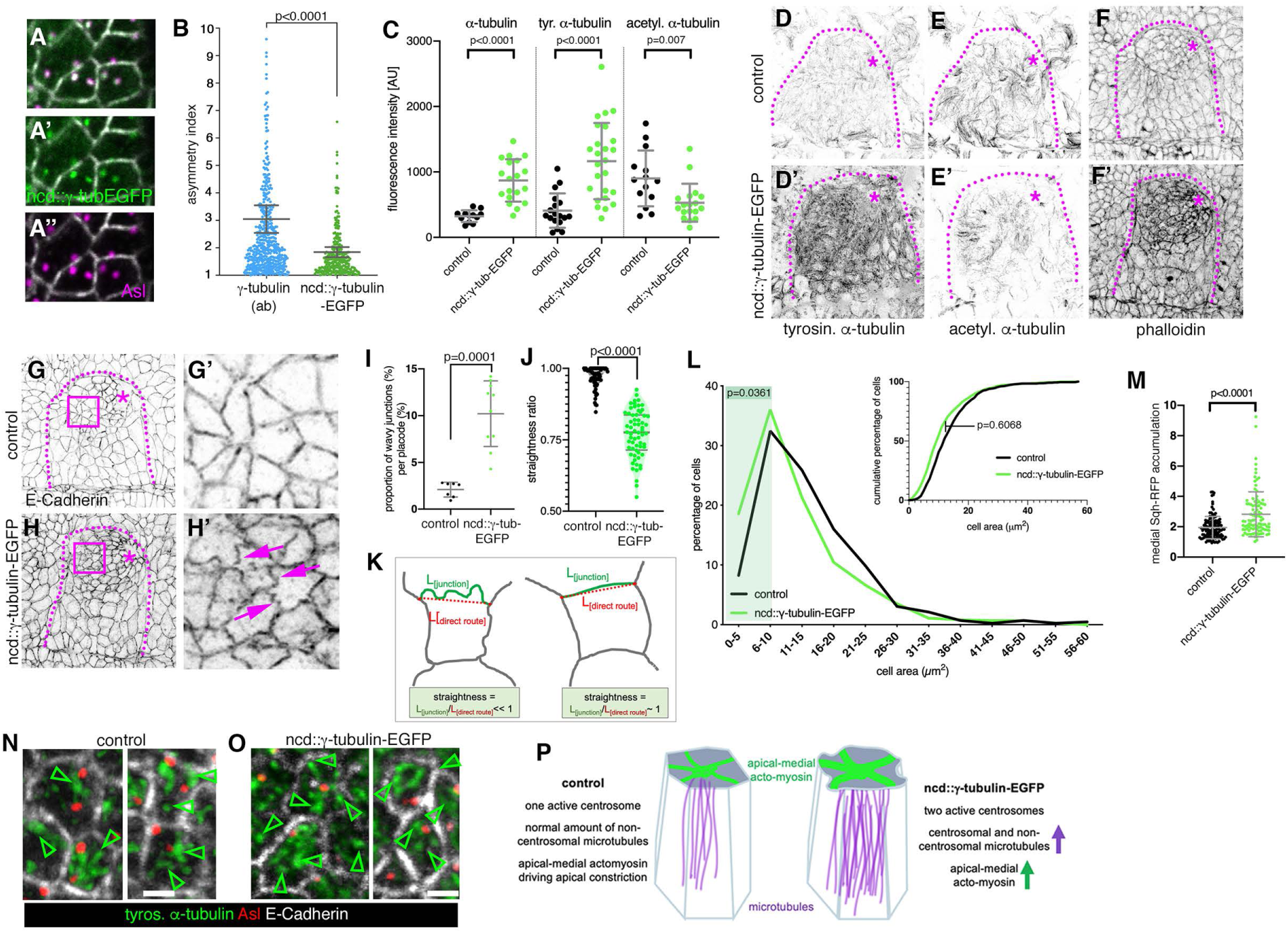
γ-tubulin overexpression increases microtubule and actomyosin levels and leads to excessive apical constriction in the placode. **A-B** Overexpression of γ-tubulin using *ncd::γ-tubulin-EGFP* leads to reduction of γ-tubulin asymmetry at centrosomes (localisation in **A-A’’** and quantification in **B**). **B** *ncd::γ-tubulin-EGFP* asymmetry was quantified as part of the data illustrated in Fig.2C.C’. The γ-tubulin (ab) data are reproduced here in comparison to the *ncd::γ-tubulin-EGFP* overexpression: with n=284 cells from 8 embryos; statistical significance was deduced by Mann-Whitney test of comparison as p<0.0001, shown are mean and 95% CI. **C** Overexpression of *ncd::γ-tubulin-EGFP* and loss of γ-tubulin asymmetry leads to increased levels of microtubule labeling using antibodies against α-tubulin or tyrosinated α-tubulin, but a slight reduction in acetylated α-tubulin labeling. Statistical significance was determined by unpaired t-tests with Welch’s correction for α-tubulin and tyrosinated α-tubulin and a Mann-Whitney test for acetylated α-tubulin with the indicated p values, shown are mean +/- SD. **D-F’** Labeling of stage 11 placodes of control and γ-tubulin overexpressing embryos, showing the increase in tyrosinated α-tubulin (**D, D’**) and phalloidin labeling to reveal F-actin (**F, F’**) and the decrease in acetylated α-tubulin labeling (**E, E’**). **G-H’** Placodes of control and γ-tubulin overexpressing embryos stained for E-Cadherin to reveal cell outlines. The magenta boxed areas in **G** and **H** are magnified in **G’** and **H’**. The dotted lines denote the placode boundary, the asterisks the invagination pit. Note the wavy junctions in placodes of *ncd::γ-tubulin-EGFP* embryos indicated by magenta arrows in **H’**. **I-K** Quantification of junction waviness. **I** Placodes of *ncd::γ-tubulin-EGFP* embryos show a significantly higher proportion of wavy junctions than control placodes (7 placodes for control and 10 placodes for *ncd::γ-tubulin-EGFP* were analysed); p= 0.0001, determined by Unpaired Mann-Whitney t-test, shown are mean +/- SD. **J** Wavy junctions in *ncd::γ-tubulin-EGFP* overexpressing embryos are significantly less straight (90 junctions from 5 placodes) than randomly picked junctions in control placodes (90 junctions from 5 placodes). Statistical significance was determined using Unpaired Mann Whitney t-test with p<0.0001, shown are violin plots with median and quartiles. **K** The straightness of a junction is defined as the ratio of the length of the junction itself (L_[junction]_) divided by the length of the direct route between vertices (L_[direct route]_) ^32, 51^. For a straight junction this value is close to 1, for a wavy junction it is <<1. **L** Apices of secretory cells of *ncd::γ-tubulin-EGFP* embryos that overexpress γ-tubulin are more constricted than apices of control placodes, illustrated are both percentage of cells of a certain apical area bin as well as the cumulative percentage of cells. 738 cells were analysed in *ncd::γ-tubulin-EGFP* embryos and 425 cells in control placodes; Kolmogorov-Smirnov two-sample test on the cumuilative data did not show a significant difference between control and γ-tubulin overexpressing embryos (p=0.6068). However, comparisons of only cells with small apices showed a significant difference, as shown when comparing the distribution of cells with apical areas between 0 and 5 μm^2^ (p=0.0361) and cells with apical areas in a range between 5 and 10 μm^2^ (p=0.0361), indicated by green shaded area. **M** *ncd::γ-tubulin-EGFP* embryos that overexpress γ-tubulin show an increase in apical-medial myosin compared to control (visualised using *sqh-RFP* embryos). 100 cells in 4 placodes were analysed in *ncd::γ-tubulin-EGFP* embryos and 125 cells in 5 control placodes; statistical significance was deduced by Mann-Whitney test as p<0.0001, shown is mean +/-SD. **N, O** Microtubules within the apical-medial region of placodal cells of *ncd::γ-tubulin-EGFP* embryos that overexpress γ-tubulin remain organised in a non-centrosomal fashion (**O**), as in the control (**N**). Arrowheads point to ends of microtubules or microtubule bundles away from centrosomes in this apical section of placodal secretory cells. Microtubules are labeled using anti tyrosinated α-tubulin (green), cell outlines are marked by E-Cadherin (white) and centrosomes by anti-Asterless (red). Scale bars are 1µm. **P** Model of the effect of γ-tubulin overexpression in placodal cells: in contrast to the control, when γ-tubulin is overexpressed both centrosomes contain similar levels of γ-tubulin and thus both nucleate microtubules. The increase in microtubule nucleation in placodal cells leads to an increase in centrosomal and also non-centrosomal microtubules, with a concomitant increase in apical-medial actomyosin and thus an increase in apical constriction.

Thus, not only did post-mitotic placodal cells show a distinctive centrosome asymmetry, similar to that previously only observed in neuroblasts, but this asymmetry was important for the establishment of a non-centrosomal microtubule array with the right amount of microtubules. Our data suggest that an excessive microtubule array, by enhancing constriction across the placode, can interfere with wild-type tube budding.

### Katanin accumulates at the apical-medial site of placodal cells

As one centrosome retained microtubule nucleation capacity, but placodal microtubules became non-centrosomally organised in constricting cells ^8^, we initially hypothesised that relocalisation of active γ-tubulin-dependent nucleation to a non-centrosomal MTOC such as the apical domain could be at play ^34^. However, we did not detect any accumulation of γ-tubulin outside of centrosomes within the placodal epidermal cells (Fig. 2B and Fig. 3A,A’). Alternatively, it has been proposed that non-centrosomal microtubules could be generated via the release of centrosomal microtubules by severing enzymes followed by selective capture and stabilisation of microtubule minus-ends ^35, 36^. One such severing enzyme is Katanin, which is a multi-subunit microtubule severing enzyme ^37^. Using a YFP-exon trap line of the regulatory subunit Katanin 80, a line tagging the endogenous locus and thus faithfully reporting expression and localisation of the endogenous protein, we discovered that Katanin 80 was selectively enriched in bright foci in the placodal cells compared to the surrounding epidermis from early stage 11 onwards (Fig. 4A-A’’). Within the placodal cells about to or undergoing apical constriction, foci of Katanin 80-YFP were localised in an apical-medial position (Fig. 4A’’). This medial localisation was close to centrosomes in just under half of all cells that showed a clear Katanin 80-YFP signal, and the localisation was more diffuse or junctional in other cells (Fig. 4B-B’). Furthermore, Katanin 80-YFP near centrosomes colocalised with the apical foci demarcating ends of microtubules bundles (Fig. 4C). In cells where Katanin localised near a single centrosome, this was the one enriched in Cnn and thus the one still nucleating microtubules (Fig. 4D,D’).

**Figure 4.**
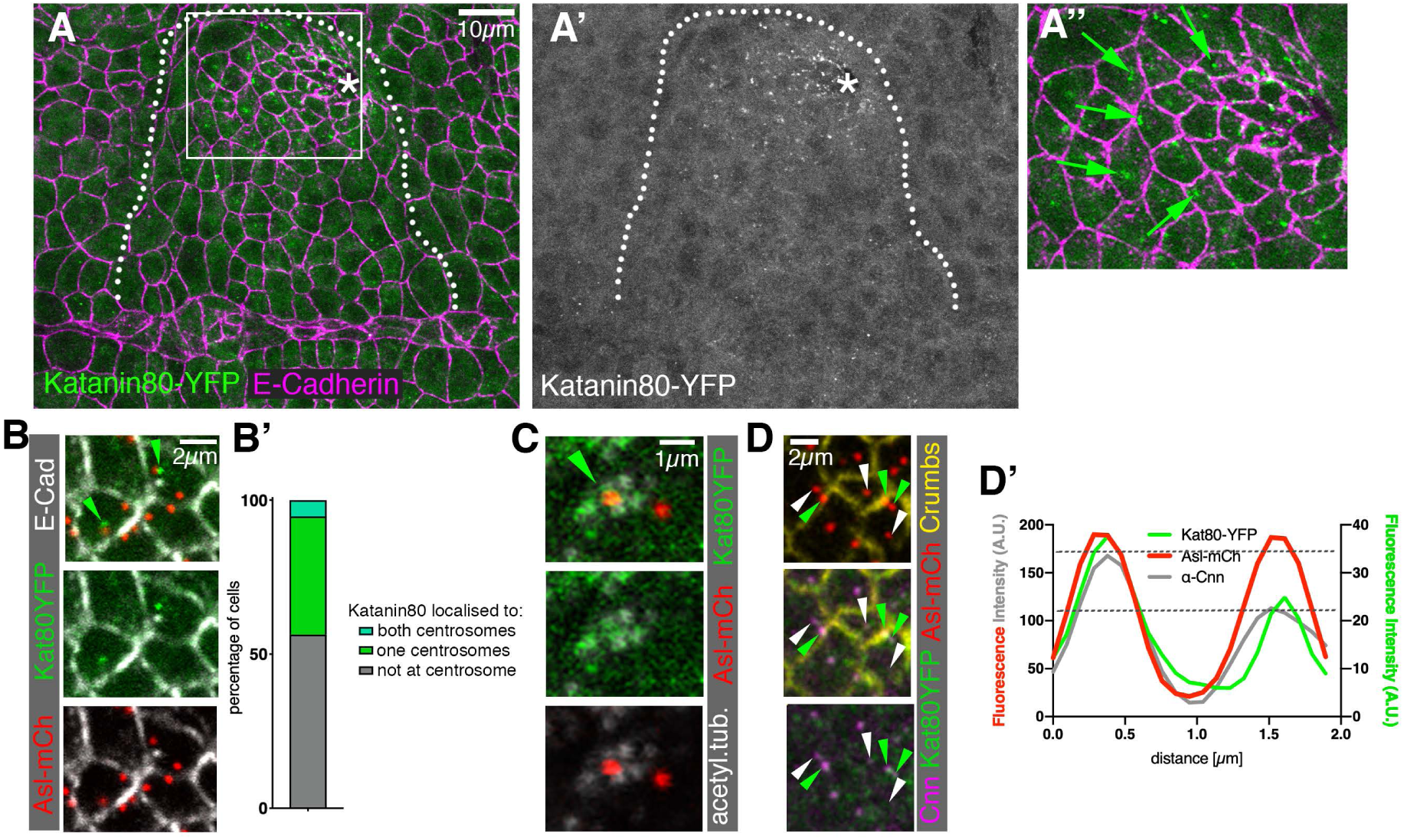
The microtubule-severing protein Katanin specifically accumulates at the apical-medial side of placodal cells. **A-A’’** Katanin80, labelled using a YFP-protein trap line (green in **A, A’’** and single channel in **A’**), accumulates specifically in the secretory placodal cells and not the surrounding epidermis. **A’’** Katanin80-YFP foci are found in an apical-medial position in the constricting population of cells (green arrows). Cell outlines are marked by E-Cadherin (magenta in **A, A’’**). The salivary gland placode is indicated by a white dotted line and the invagination point by an asterisk. **B-C** Katanin80-YFP accumulates near centrosomes and microtubules. **B** Kat80YFP in green localises close to centrosomes marked by Asl-mCherry in red, cell outlines in white are labeled by E-Cadherin. **B’** Quantification of Katanin80-YFP accumulation near centrosomes. **C** Katanin80-YFP (green) can be found near microtubules labelled by staining for acetylated α-tubulin (white) in the proximity of centrosomes marked by Asl-mCherry (red). **D-D’** Katanin80-YFP (green) accumulates near the microtubule-nucleating centrosomes marked by Cnn (magenta), with all centrosomes marked by Asl-mCherry (red) and cell outlines marked by Crumbs (yellow). **D’** Line scan profile through both centrosomes of a single cell to illustrate the co-enrichment of Katanin80-YFP on the Cnn-enriched centrosome. Centrosome positions are marked by Asl-mCherry.

Thus, Katanin’s expression and localisation in placodal cells strongly suggested that it plays a role in the generation of the non-centrosomal microtubule array.

### Loss of Katanin disrupts the non-centrosomal microtubule array and apical-medial actomyosin-mediated apical constriction

In order to test whether Katanin-severing was important for the generation of the non-centrosomal microtubule array within the placodal cells and for tube budding morphogenesis, we selectively depleted Katanin 80-YFP from the placodal cells using the degradFP system (Fig. 5A). This system can target GFP-and YFP-tagged endogenous proteins for degradation by the proteasome through tissue-specific expression of a modified F-box protein that is fused to an anti-GFP nanobody (^38^; Fig.5A). Expression of degradFP specifically in the salivary gland placode under *fkhGal4* control led to a reduction of Katanin80-YFP levels in the placodal cells (Fig. 5B-D). In contrast to the control, in these Katanin-depleted cells microtubules did not lose contact with centrosomes but rather remained in close association with the centrosomes labeled by Asl. This led to microtubule bundles being visible running parallel to the apical domain, rather than the foci of microtubule minus ends being visible as in the control (Fig. 5E-G).

**Figure 5.**
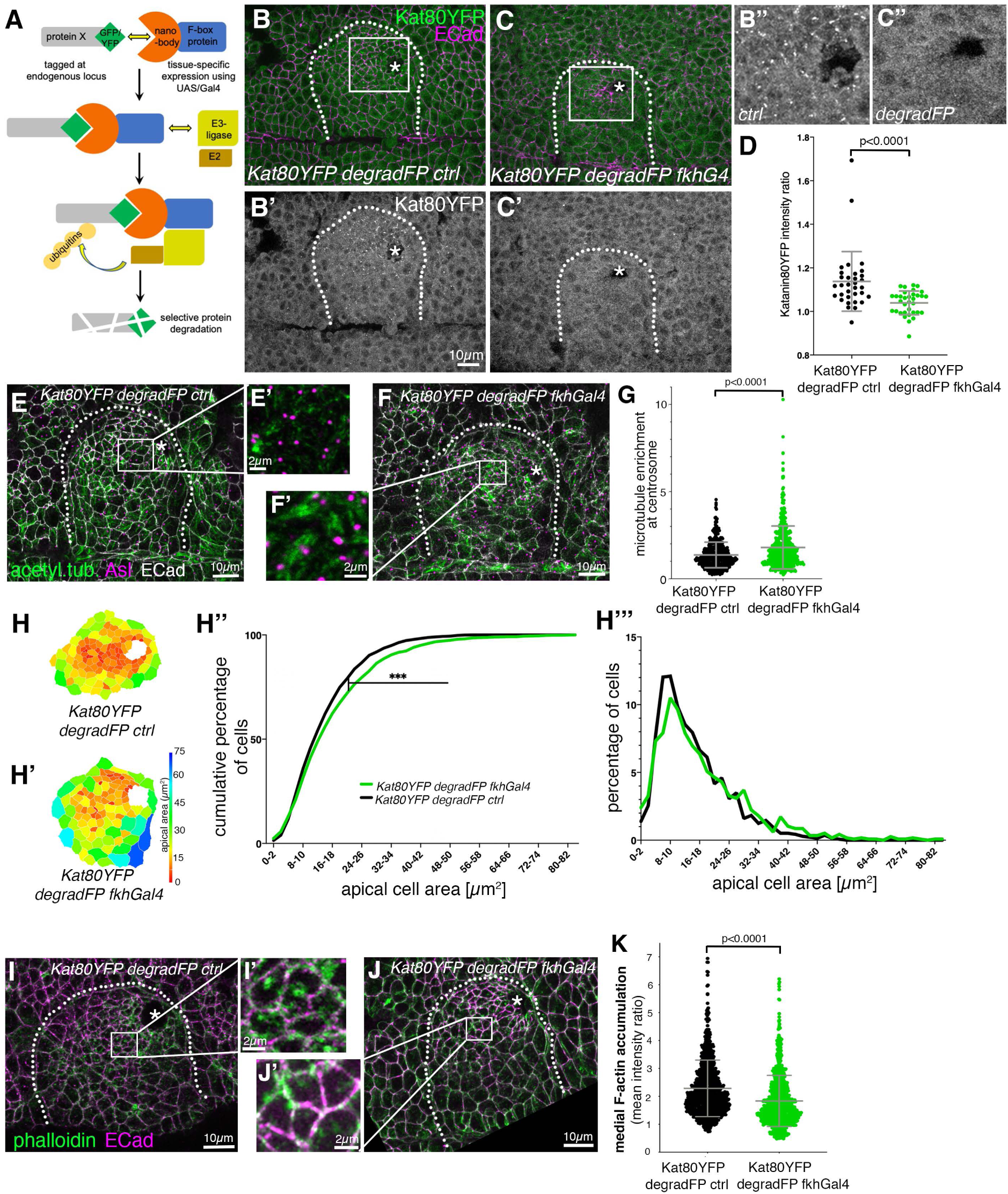
Loss of Katanin disrupts the non-centrosomal microtubule array and apical-medial actomyosin-mediated apical constriction. **A** Schematic of the ‘degradFP’ tissue-specific-degradation system (as in ^38^): tissue-specific expression of an F-box-anti/GFP-nanobody fusion protein, degradFP, (using UAS/Gal4; ^52^) leads to tissue-specific degradation of any endogenously GFP/YFP-tagged protein, in this case Katanin80-YFP. **B-D** Expressing degradFP using *fkh-Gal4* in the salivary gland placode (*Kat80YFP degradFP fkhG4*; **C-C’’**) leads to significant loss of Katanin80-YFP compared to control (*Kat80YFP degradFP ctrl*; **B-B’’**). Cell outlines are marked by E-Cadherin (E-Cad). **D** Quantification of Katanin80-YFP depletion (*Kat80YFP degradFP ctrl* n=33; *Kat80YFP degradFP fkhG4* n=34/ shown are mean +/-SD, statistical significance was determined by Mann-Whitney test as p<0.0001). **B’’** and **C’’** are higher magnifications of the white boxes marked in **B** and **C**, respectively. **E-G** In placodes where Katanin80-YFP is degraded (**F, F’**), microtubules (green, labeled for acetylated α-tubulin) remain localised within the apical domain and in contact with centrosomes (magenta, labeled for Asl) compared to control (**E, E’**) where a non-centrosomal longitudinal array is formed. **E’** and **F’** are magnifications of the areas indicated in **E** and **F** by a white box. **G** Quantification of the effect shown in **E-F** (*Kat80YFP degradFP ctrl*: 440 cells from 14 embryos; *Kat80YFP degradFP fkhG4*: 547 cells from 20 embryos; shown are mean +/-SD, statistical significance was determined by Mann-Whitney test as p<0.0001). **H-H’’’** Katanin80-YFP degradation (**H’**) leads to a loss of apical constriction compared to control (**H**), apical area of cells of example placodes are shown in a heat map. **H’’** Quantification of apical area distribution of placodal cells in control (WT) and Katanin depleted (degradFP) placodes at stage 11 showing the cumulative percentage of cells relative to apical area size. **H’’’** Percentage of cells in different size-bins [Kolmogorov-Smirnov two-sample test, P << 0.001 (***)]. 10 placodes were segmented and analysed for each condition, the total number of cells traced was N(*Kat80YFP degradFP ctrl*)=1373, N(*Kat80YFP degradFP fkhG4*)=1162. **I-K** In placodes where Katanin80-YFP is degraded (**J, J’**), apical-medial F-actin (green, labeled using phalloidin) is reduced compared to control (**I, I’**) where apical-medial actin is highly prevalent. **I’** and **J’** are magnifications of the areas indicated in **J** by a white box. **K** Quantification of loss of apical-medial F-actin (*Kat80YFP degradFP ctrl:* 924 cells from 17 embryos; *Kat80YFP degradFP fkhG4:* 921 cells from 15 embryos/shown are mean +/-SD, statistical significance was determined by Mann-Whitney test as p<0.0001). The salivary gland placode is indicated by a white dotted line and the invagination point, where present, by an asterisk.

We have previously shown that overall loss of microtubules in the placodal cells led to a failure of apical constriction of the cells during budding, due to a selective loss of apical-medial actomyosin ^8^. We therefore analysed whether failure in the generation of the longitudinal non-centrosomal array and continued centrosomal anchoring affected apical constriction. We quantified the apical cell area of placodal cells at a timepoint where the invagination pit had formed (mid-to-late stage 11; Fig. 5H-H’’’). Though apical constriction was not abolished, there was a significant reduction in the amount of apically constricted cells when Katanin was degraded (Fig. 5H’’-H’’’). Furthermore, analysis of apical actin accumulation revealed that apical-medial actin levels were reduced (Fig. 5I-K).

Thus, Katanin-mediated microtubule severing at a single active centrosome plays a key role in the formation of the longitudinal non-centrosomal microtubule array that in turn supports apical constriction.

### Patronin localises specifically to non-centrosomal microtubule minus ends in apical-medial region of placodal cells

Free microtubule minus ends generated by Katanin severing activity tend to be highly susceptible to depolymerisation and usually require stabilisation to prevent this happening ^39, 40^. Members of the family of CAMSAPs (<u>Ca</u>l<u>m</u>odulin-regulated-<u>S</u>pectrin-<u>a</u>ssociated <u>p</u>roteins), with the single orthologue in flies called Patronin, have such a capacity for minus-end stabilisation and also anchoring at non-centrosomal sites ^41, 42^. Whereas endogenously tagged Patronin (Patronin-YFP; ^18^) accumulated near adherens junctions in most epidermal cells outside the placode (Fig. 6A-C’), consistent with a previously described function in stabilising microtubules in this location ^43^, within the placodal cells undergoing apical constriction Patronin accumulated in an apical-medial position (Fig. 6A-B’). Just upon salivary gland placode specification at the end of stage 10, but prior to the microtubule rearrangement, Patronin was also localised predominantly to adherens junctions in placodal cell (Fig. S4A-C), but from early stage 11 onwards it was strongly localised in the apical-medial position, where it now colocalised with microtubule minus ends (Fig. 6D-E; ^8^). This dynamic relocalisation of Patronin accumulation depended on an intact microtubule cytoskeleton, as its apical-medial localisation in the placodal cells reverted to a junctional localisation upon depletion of microtubules via Spastin overexpression within the salivary gland placode only (using *fkhGal4*; (Fig. 6F-H)). Furthermore, the recruitment of Patronin also depended on the action of Katanin: when Katanin 80-YFP was degraded in the salivary gland placode using *fkh-Gal4 UAS-degradFP*, a Patronin-RFP transgene was mainly localised at apical junctions (Fig. 6J’-K), whereas in the control it accumulated in the apical-medial position (Fig. 6I-I’’ and K).

**Figure 6.**
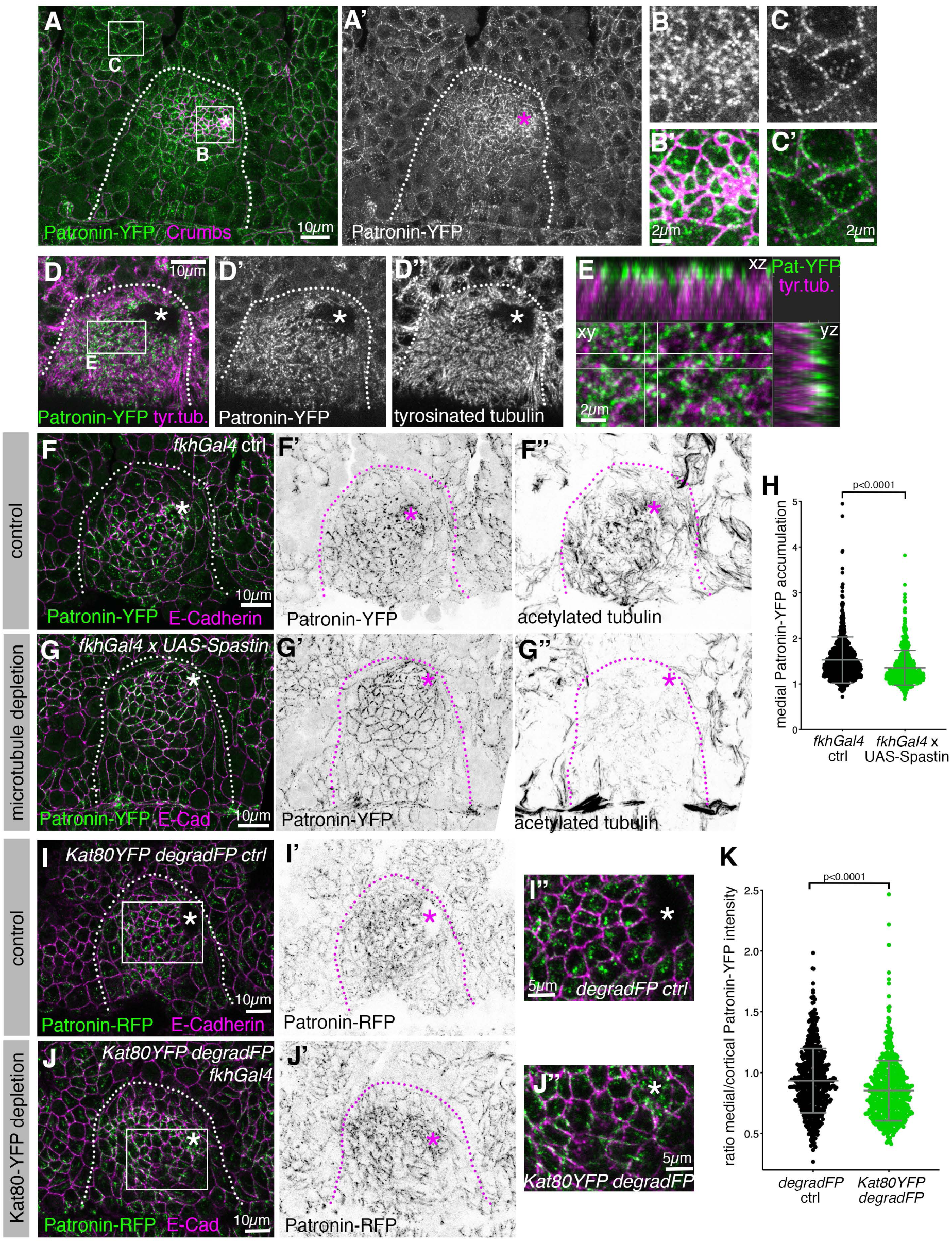
Patronin localises specifically to non-centrosomal microtubule minus ends in the apical-medial region of placodal cells. **A-C’** Patronin, labeled using an endogenously labeleld YFP-tagged protein trap line, Patronin-YFP (green and **A’, B, C**), accumulates at adherens junctions throughout the embryonic epidermis (**A** and **C, C’**), but in the apically-constricting cells of the salivary gland placode accumulates in an apical-medial position (**A** and **B, B’**). Junctions are labelled using Crumbs (magenta). **B-C’** are magnification of the white boxes in **A**. **D-E** Apical-medial Patronin-YFP (green and **D’**) localises to the ends of longitudinal microtubules labeled for tyrosinated α-tubulin (magenta and **D’’**). **E** is a magnification of the white box in **D** and also shows corresponding xz and yz-cross-sections. **F-H** Patronin depends on longitudinal microtubules for its apical-medial localisation. In contrast to control (**F-F’’**) where Patronin-YFP (green and **F’**) localises to apical-medial sites, when microtubules are lost upon expression of *UAS-Spastin* under *fkh-Gal4* control (**G-G’’**), Patronin-YFP (green and **G’**) continues to be localised to junctional sites and does not relocalise to apical-medial sites. Junctions are labelled for E-Cadherin (magenta), and microtubules are labeled for acetylated α-tubulin (**F’’, G’’**). **H** Quantification of reduction of apical-medial Patronin-YFP upon placodal microtubule loss (*fkhGal4 ctrl:* 577 cells from 13 embryos; *fkhGal4 x UAS-Spastin:* 955 cells from 19 embryos/shown are mean +/- SD, statistical significance was determined by Mann-Whitney test as p<0.0001). **I-K** Depletion of Katanin80-YFP using *degradFP x fkhGal4* (**J-J’’**) leads to loss of apical-medial Patronin-RFP (green and **I’, J’**) in contrast to control (**I-I’’**). **I’’** and **J’’** are magnifications of the white boxes shown in **I** and **J**. Membranes are labelled by E-Cadherin (magenta). **K** Quantification of changes of the apical-medial vs apical-junctional Patronin-RFP localisation upon degradation of Katanin80-YFP (*Kat80-YFP degradFP ctrl:* 600 cells from 12 embryos; *Kat80-YFP degradFP fkhGal4:* 595 cells from 12 embryos/shown are mean +/- SD, statistical significance was determined by Mann-Whitney test as p<0.0001). In overview panels the salivary gland placode is indicated by a dotted line and the invagination point by an asterisk.

Thus, we identified Patronin as the factor that binds the minus ends of the longitudinal microtubule array in placodal cells, once microtubules were released by Katanin from the nucleating centrosome.

### Loss of Patronin function prevents the proper organisation of the non-centrosomal microtubule array in the placode

Patronin is required during oogenesis ^18^ and therefore we could not generate maternal/zygotic mutants that would lack all Patronin protein in the embryo. Zygotic loss of Patronin alone only led to weakly penetrant phenotypes in salivary gland tube budding, most likely due to a rescue by perdurance of maternal RNA and protein (data not shown). Also, Patronin in the apical-medial position of placodal cells bound to microtubule minus ends appeared to be very stable as several approaches, such as targeting Patronin by RNAi (Fig. 7) or tissue-specific degradation using a degradFP approach (Fig. S5), only led to a reduction in Patronin protein levels but not a complete loss (Fig. 7 and Fig. S5). To test Patronin requirement and reduce its levels, we used expression of an RNAi construct against *patronin* mRNA (*UAS-Patronin-RNAi*) under the control of a ubiquitous embryonic driver, *daughterless-Gal4, DaGal4*, (Fig. 7), thereby also bypassing an earlier requirement for Patronin during gastrulation ^6^. In placodes of *UAS-Patronin-RNAi x DaGal4* embryos (Fig. 7A-E’) medial Patronin fluorescence intensity was reduced by 24.4% (Fig. 7E), whereas junctional Patronin intensity in placodes was reduced only by 14.9% (Fig. 7E’). Nonetheless, although fluorescence intensity of acetylated α-tubulin staining was not overall reduced within the whole apical domain of placodal cells (Fig. S5C), there was a clear change in organisation of the microtubule array (Fig. 7A’’, B’’, C, D). At stage 11 in the control, apical foci of longitudinal microtubule bundles were visible within the apical domain of cells close to the invagination point (Fig. 7C), colocalising with medial Patronin foci (Fig. 7C’). Instead, when medial Patronin was lost due to Patronin-RNAi, microtubules were observed lying within the apical domain, often terminating in close proximity to junctions (Fig. 7D), where Patronin was still localised (Fig. 7D’). Placodes of *UAS-Patronin-RNAi x DaGal4* embryos often showed a disrupted morphology, and apical constriction appeared less efficient (Fig. 7F-H’), most likely because Patronin protecting the longitudinal arrangement of microtubules was important for the role of microtubules in supporting apical-medial actomyosin that we demonstrated previously ^8^. This is also supported by the observation that medial Patronin foci in cells near the invagination point are dynamic and coalesce during an apical constriction pulse (Fig. S4D-F), as also seen during gastrulation ^6^.

**Figure 7.**
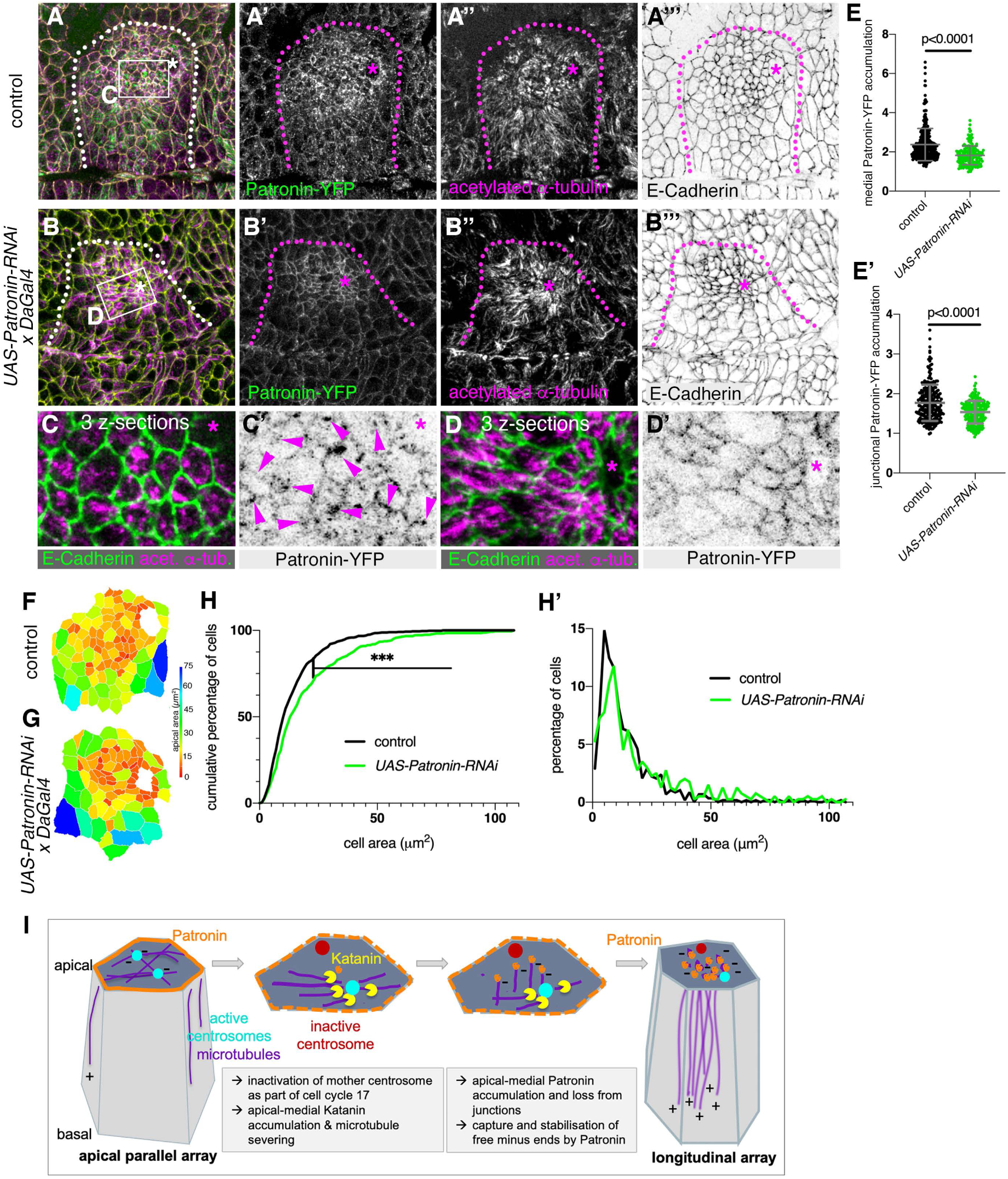
Loss of Patronin affects microtubule organisation and apical constriction in the placode. **A-D’** Patronin (**A’, B’, C’, D’**) and acetylated microtubule (**A’’, B’’** and magenta in **C, D**) labeling in control embryos (**A-A’’’** and **C**,**C’**) and *UAS-Patronin-RNAi x DaGal4* embryos (**B-B’’’** and **D, D’**). **E-E’** Quantification of medial (**E**) and junctional (**E’**) Patronin fluorescence intensity in control and *UAS-Patronin-RNAi x DaGal4* embryos. 250 cells from 6 placodes were analysed in the control and 214 cells from 5 placodes for *UAS-Patronin-RNAi x DaGal4* embryos; shown are mean +/- SD, statistical significance was determined by Mann-Whitney test as p<0.0001. **F-H’** Depletion of Patronin in *UAS-Patronin-RNAi x DaGal4* embryos (**G**) leads to a loss of apical constriction compared to control (**F**), apical area of cells of example placodes are shown in a heat map. **H, H’** Quantification of apical area distribution of placodal cells in control and Patronin-depleted (*UAS-Patronin-RNAi x DaGal4*) placodes at stage 11 showing the cumulative percentage of cells relative to apical area size (**H**) as well as the percentage of cells in different size-bins (**H’**) [Kolmogorov-Smirnov two-sample test, P << 0.001 (***)]. 6 placodes were segmented and analysed for each condition, the total number of cells traced was N(*control*)=585, N(*UAS-Patronin-RNAi x DaGal4*)=399. **I** Model of generation of the longitudinal non-centrosomal microtubule array in salivary gland placodal cell prior to apical constriction: inactivation of the mother centrosome concomitant with entering cycle 17 and becoming post-mitotic leads to restriction of microtubule nucleation to a single centrosome. An increase in apical-medial Katanin levels in the secretory cells drive severing of microtubules at the centrosome. Severed microtubule minus ends are then captured by apical Patronin in the apical-medial region, thereby promoting the longitudinal microtubule arrangement.

Thus, Patronin in the salivary gland placode, once localised to microtubule minus ends within the apical-medial domain, serves to support the organisation of the longitudinal microtubules array that in turn is required for successful apical constriction of cells and formation of the wild-type tubular organ.

## Discussion

The organisation of the microtubule cytoskeleton is key to many cellular functions, both in individual cells as well as cells in the context of a tissue. In many actively cycling cells the interphase microtubule cytoskeleton is organised from centrosomes as the major MTOCs. In contrast, in many differentiated cells microtubules are nucleated or anchored from sites independent of centrosomes to support specific cellular functions. This is true for most epithelia and neurons in animals but also yeast and plant cells.

Despite non-centrosomal microtubule arrays being common features in such differentiated cells, the mechanisms of their generation and organisation are still unclear ^35^. On the one hand, non-centrosomal microtubules can be *de novo* nucleated at non-centrosomal MTOCs, and for this process γ-tubulin as part of the γ-TURC is essential ^12, 34^. However, the mechanisms leading to recruitment of γ-tubulin at non-centrosomal MTOCs is still poorly understood. For example, a splice variant of Cnn, CnnT, resides at mitochondria in *Drosophila* spermatids to recruit γ-tubulin and convert the mitochondria into MTOCs ^44^. However, *de novo* nucleation at non-centrosomal MTOCs is only one way of generating a non-centrosomal array. Rather than relocalising the nucleation capacity away from centrosomes, a release and capture of microtubules generated at nucleating centrosomes is an alternative mechanism to form non-centrosomal microtubule arrays that has been first observed in neurons ^45^. Furthermore, once even a small cluster of non-centrosomal microtubules has formed, this organisation can be selectively amplified by targeted severing of such microtubules combined with capture of newly generated minus ends and continued polymerisation from free plus ends.

Tube budding morphogenesis of the salivary glands in the *Drosophila* embryo, and in particular the apical constriction of cells leading to tissue bending, depends on an intact microtubule cytoskeleton that is organised as a non-centrosomal longitudinal array ^8^. Here we elucidate the mechanism by which this non-centrosomal microtubule array is formed. Interestingly, the initial formation of the array still involves centrosomal nucleation, though in the salivary gland placodal cells this is restricted to the daughter centrosome of the last division that retains nucleation capacity. Both non-centrosomal array formation as well as the downstream morphogenetic process of apical constriction also require the action of the severing enzyme Katanin, as well as the minus-end stabiliser Patronin ^37, 41^. Thus, in this tissue a mechanism appears to operate whereby at least initially centrosomally-nucleated microtubules are severed by Katanin at the centrosome and their free minus ends then recruit Patronin. Minus ends remain anchored within the apical-medial region ^8^ and further research will elucidate whether Patronin also interacts with other potential binding partners that are localised in this region, such as the spectraplakin Shot ^8^.

It is curious to speculate what the close apposition and interaction of microtubules, and in particular their minus ends, and apical-medial actomyosin in apically-constricting cells entails. Our past and recent data strongly suggest a regulatory interplay, with the presence and amount of non-centrosomal microtubules directly affecting presence and amount of apical-medial actomyosin and thus the rate of apical constriction in placodal cells. The loss of microtubules leads to loss of apical-medial myosin and hence reduction in apical constriction ^8^, and increase in placodal non-centrosomal microtubules leads to an increase in apical-medial actomyosin and increased apical constriction, as described above. Many questions remain, such as whether this interaction is regulatory in that microtubule minus ends near apical-medial actomyosin could serve to guide directional transport of vesicles to the apical domain, and recent evidence suggests that this might at least partially be the case ^46^. There could be a mechanical requirement for a close apposition of longitudinal microtubules and apical-medial actomyosin in allowing the formation of a wedged shape of an apically constricting cell, akin to ideas about cellular ‘tensegrity’ originally proposed by Ingber et al. ^47^. How the latter could be tested experimentally is not clear, though *in silico* modelling might pave a way for a better understanding of mechanical implications of this apposition.

Interestingly, a different requirement for microtubules in assisting actomyosin-based apical constriction was recently reported during mesoderm invagination in the *Drosophila* embryo. Here, disrupting microtubules through colchicine or taxol injections to either depolymerise or stabilise the network acutely led to longer persistence and size of the interconnected apical-medial actomyosin network that stretches across cells ^6^. Microtubules appear necessary to induce the actin turnover near junctions where the apical-medial foci connect via cell-cell adhesions between neighbouring cells.

Also in the early embryo during gastrulation, Patronin appears to behave more dynamically and affected microtubules in a different way, suggesting that roles for Patronin might be developmental-stage specific. Expression of shRNA targeting Patronin at this early stage led to a strong overall loss of Patronin, and a concomitant loss of acetylated microtubules ^6^. Several hours further into embryogenesis, during the formation of the salivary gland tubes analysed here, Patronin is more stable and resistant to depletion (Fig. 7 and S5) and its loss or reduction does not lead to concomitant loss but rather disorganisation of microtubules. Within the placode, the reduction in apical-medial Patronin without overall loss of microtubules could be due to junctional Patronin now binding microtubules that were severed at centrosomes and require anchoring. Alternatively, as has been described in cells in culture, loss or reduction of Patronin/CAMSAPs can directly affect the ability of Katanin to sever microtubules ^48^, and thus upon Patronin depletion more microtubules may remain attached to the nucleating centrosome. Supporting this, Patronin colocalises with a pool of Katanin in the secretory cells of the placode (Fig. S5). Such a disorganisation of microtubules upon perturbance of Patronin is also more reflective of its behaviour in the adult epithelium of the follicle cells in the fly ovary ^49^ and the small intestine of postnatal mice ^9^. This most likely reflects changes from actively dividing to post-mitotic epithelial behaviour in both tissues, and is also supported by observed changes to CAMSAP3 mobility during maturation of Caco2 cell cysts in culture ^50^.

Thus, it appears that depending on the tissue context, including the mitotic activity that epithelial cells display at a given time, the interplay of centrosome-nucleated microtubules, microtubule-release from centrosomes as well as the function of CAMSAP-family proteins such as Patronin are highly coordinated and adjusted to serve the assembly and maintenance of particular microtubule arrays. In the case of the formation of the tubular organ of the salivary glands, a key process of tube formation, we have elucidated a key mechanism that harvests the changes at centrosomes due to the cells becoming post-mitotic (Fig. 7I) and pairs it with the activity of Katanin and Patronin to promote formation and maintenance of the longitudinal non-centrosomal microtubule array that itself supports apical constriction of cells. It will be interesting to determine in the future how conserved this interplay is in other tissues reaching post-mitotic state but still undergoing morphogenetic processes such as apical constriction and tissue bending.

## Acknowledgments

The authors would like to thank the following people; for reagents and fly stocks: Debbie Andrew, Jordan Raff, Paul Conduit, Daniel St. Johnston, Claudio Sunkel, Yu-Chiun Wang, Jens Januschke, Cayetano Gonzales, Magali Suzanne, Markus Affolter, Ron Vale. The work is supported by the Medical Research Council (file reference number U105178780).

## Author contribution

Conceptualisation, K.R. and G.G.; Methodology, K.R., G.G. G.G.; Investigation, K.R., G.G., G.G.; Writing-Original Draft, K.R., G.G.; Funding Acquisition, K.R.

## Supplemental Material

**Figure S1, related to Figure 1.**
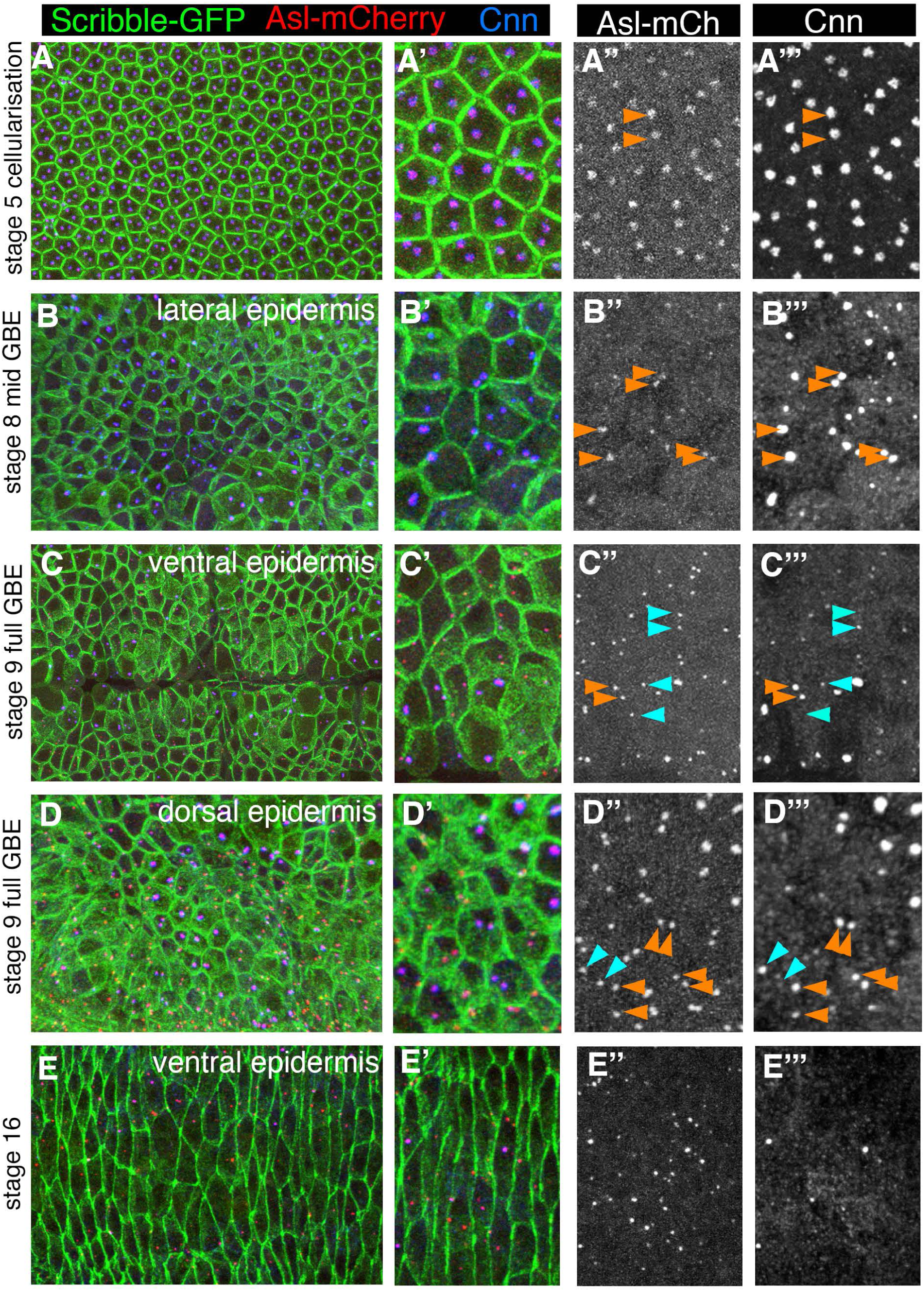
Changes to centrosomes during post-blastoderm embryogenesis. Comparison of Asl-mCherry (red, single channel in **A’’-E’’**) and Cnn (blue, single channel in **A’’’-E’’’**) levels at embryonic centrosomes at different stages and in different tissues. Cell outlines are labelled using Scribble-GFP that localises along the lateral membranes of all epithelial cells. Orange arrowheads point to centrosomes within a cell with even Cnn levels at both centrosomes, blue arrowheads point to asymmetric Cnn levels within a single cell. **A-A’’’** During and just after cellularisation at stage 5, embryonic cells contain two centrosomes with even levels of Cnn. **B-B’’’** During stage 8 in the lateral epidermis undergoing germ band extension, Cnn levels are even on both centrosomes and only strongly increased on centrosomes of cells about to undergo mitosis. **C-D’’’** At stage 9 in the ventral (**C-C’’’**) and dorsal (**D-D’’’**) epidermis, cells with asymmetric Cnn levels can be found, likely representing cells in G2 of cycle 15 or 16. These cells will divide again and thus are on the way to generating two centrosomes with equal Cnn levels. **E-E’’’** At stage 16 in the dorsal epidermis most centrosomes have lost all Cnn.

**Figure S2, related to Figure 2.**
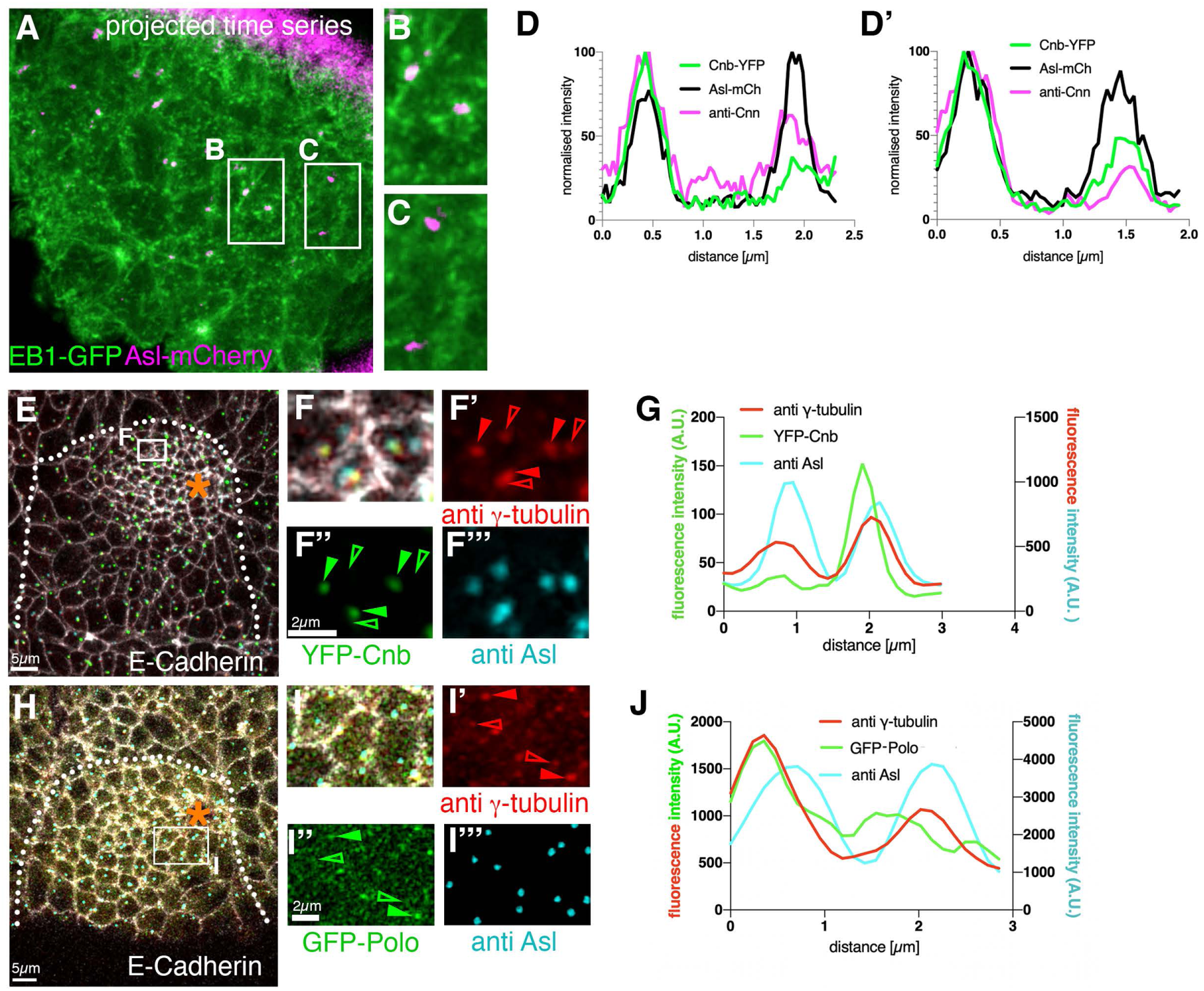
Centrosome composition and nucleation asymmetry in placodal cells. **A-C** Projected time series of *EB1-GFP Asl-mCherry* embryos in a region of a stage 11 salivary gland placode to reveal EB1-GFP ‘comets’ where they emanate from centrosomes. **B** shows a region with two Asl-mCherry centrosomes with EB1-GFP comets and thus actively nucleating microtubules, whereas **C** shows two Asl-mCherry centrosomes without comets and thus not nucleating. **D-D’** Two further examples of centrosome asymmetries of Cnn and Cnb in individual placodal cells at stage 11, as shown in Fig.2G. **E-G** Comparative analysis of YFP-Centrobin (YFP-Cnb) and γ-tubulin asymmetries at centrosomes. **E-F’’’** Overview (**E**) and higher magnifications (**F-F’’’**) of a placode, with anti E-Cadherin in white, anti γ-tubulin in red, anti Asl in cyan and YFP-Cnb in green. Filled arrowheads in **F’-F’’’** point to centrosomes with γ-tubulin and YFP-Cnb enriched, hollow arrowheads point to the centrosomes in the same cell with lower levels of γ-tubulin and YFP-Cnb. **G** Line scan profile through both centrosomes of a single cell to illustrate the co-enrichment of γ-tubulin and Cnb on the same centrosome. **H-J** Comparative analysis of GFP-Polo and γ-tubulin asymmetries at centrosomes. **H-I’’’** Overview (**H**) and higher magnifications (**I-I’’’**) of a placode, with anti E-Cadherin in white, anti γ-tubulin in red, anti Asl in cyan and GFP-Polo in green. Filled arrowheads in **I’-I’’’** point to centrosomes with γ-tubulin and GFP-Polo enriched, hollow arrowheads point to the centrosomes in the same cell with lower levels of γ-tubulin and GFP-Polo. **J** Line scan profile through both centrosomes of a single cell to illustrate the co-enrichment of γ-tubulin and Polo on the same centrosomes. The salivary gland placode is indicated by a dotted line and the invagination point by an asterisk.

**Figure S3, related to Figure 3.**
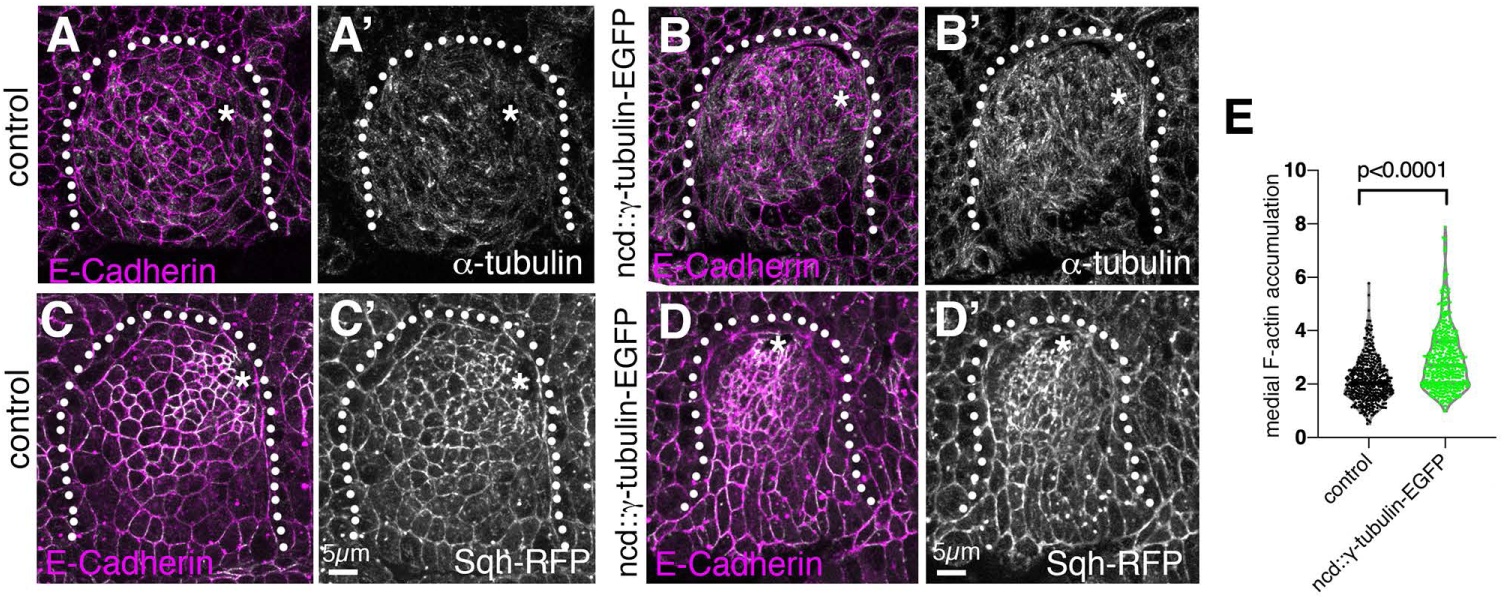
γ-tubulin overexpression increases microtubule and actomyosin levels and leads to excessive apical constriction in the placode. **A-B’** Stage 11 placode labelled for α-tubulin in a control (**A, A’**) or γ-tubulin overexpressing embryo (**B, B’**; *ncd::γ-tubulin-EGFP*). Cell outlines are marked by E-Cadherin labeling (magenta in **A, B**). **C-D’** Stage 11 placode visualising myosin using *sqh-RFP* in a control (**C, C’**) or γ-tubulin overexpressing embryo (**D, D’**; *ncd::γ-tubulin-EGFP*). Cell outlines are marked by E-Cadherin labeling (magenta in **C, D**). **E** Quantification of F-actin fluorescence in stage 11 placodes (as shown in Fig.3F.F’) in control embryos and γ-tubulin overexpressing embryos, with 480 cells from 12 embryos quantified in the control and 520 cell from 13 embryos in *ncd::γ-tubulin-EGFP* embryos. Data are shown as a violin plot with median and quartiles, statistical significance was determined by Mann-Whitney test as p<0.0001.

**Figure S4, related to Figure 6.**
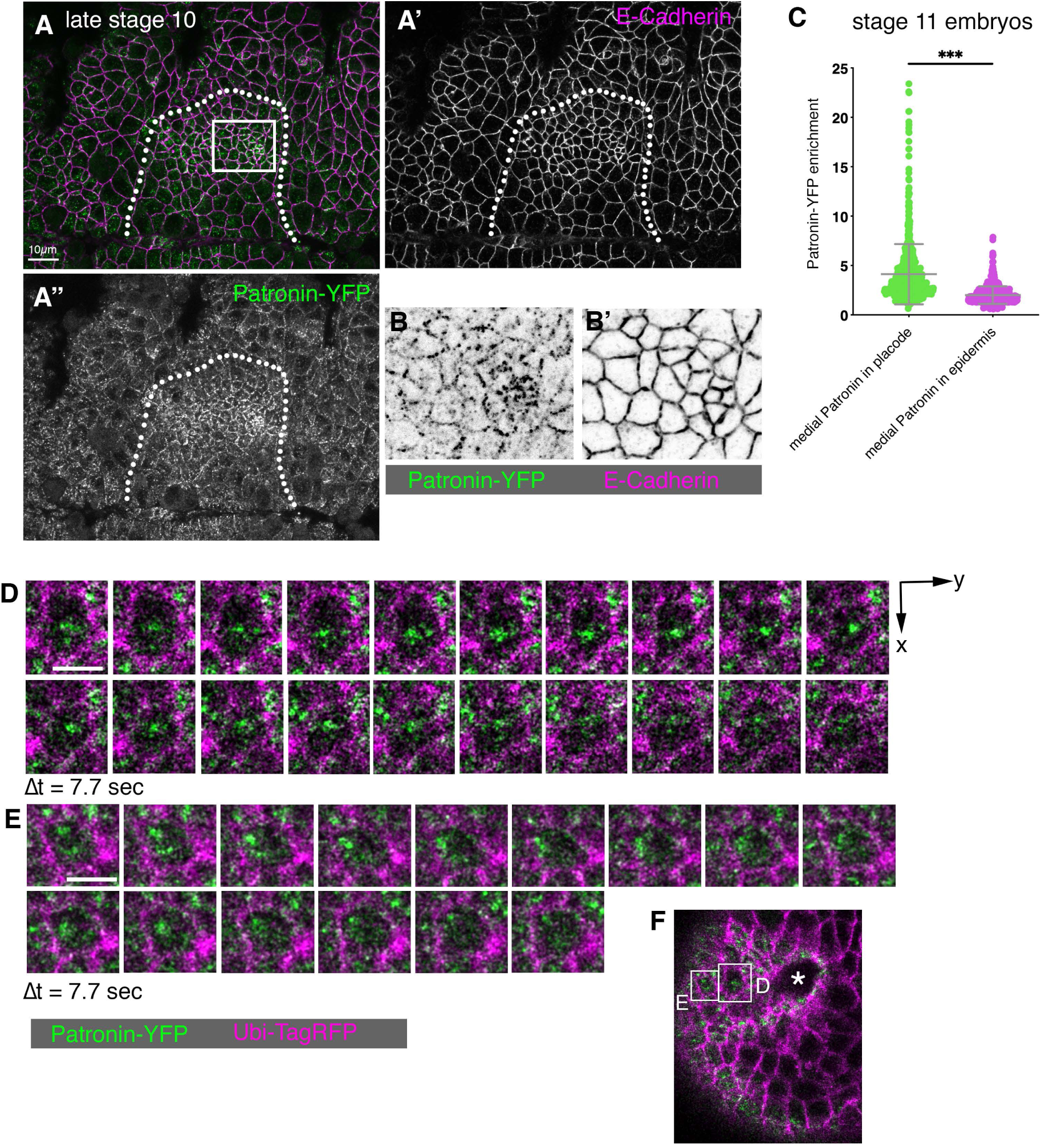
Patronin localisation and dynamics in stage 10-11 embryos. **A-B’** In a salivary gland placode at stage 10, prior to tissue bending and tube invagination commencing, Patronin-YFP localisation (green in **A**, single channel in **A’’** and **B**) is mostly restricted to adherens junctions marked by E-Cadherin (magenta in **A**, single channel in **A’** and **B’**). **C** Quantification of medial Patronin localisation at stage 11, once tube invagination has commenced, in secretory cells in the placode compared to surrounding epidermal cells. 726 cells from 15 embryos in total were analysed for the placode and epidermis, shown are mean +/-SD, statistical significance was determined using Mann-Whitney test as p<0.001 (***). **D-F** Live time-lapse analysis of Patronin-YFP; Ubi-TagRFP in a stage 11 placode in cells close to the invagination point (position of cells in **D, E** is indicated in **F** where the invagination pit is marked by an asterisk); Patronin-YFP is in green and Ubi-TagRFP in magenta. The time interval between frames (Δt) is 7.7 sec, the scale bars denote 5µm. Note the dynamic behaviour of Patronin-YFP, both in localisation as well as absolute levels.

**Figure S5, related to Figure 7.**
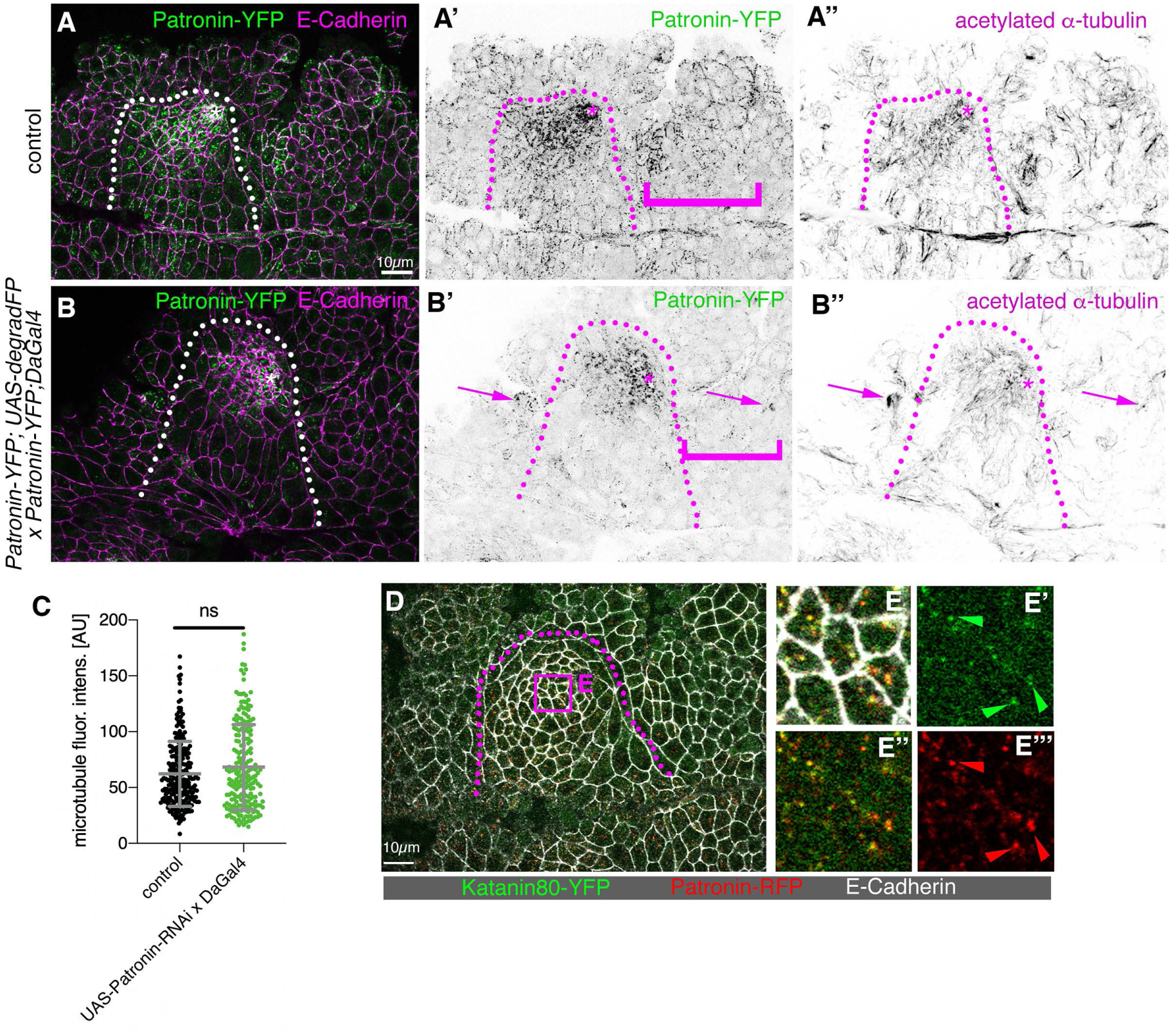
Patronin depletion and its effects. **A-B’’** Attempted Patronin depletion using degradation of Patronin-YFP via the degradFP system. In control embryos (**A-A’’**) Patronin is localised to junctions and the apical-medial region within the placode and to junctional regions in the surrounding epidermis (magenta bracket in **A’**). I n contrast in embryos of the genotype *Patronin-YFP; UAS-degradFP x Patronin-YFP; DaGal4* (**B-B’’**) most junctional Patronin-YFP has been degraded (bracket in **B’**), but much apical-medial Patronin-YFP is left in the placode and also at time in select locations in the surrounding epidermis where Patronin colocalises with acetylated α-tubulin that remains (arrows in **B’, B’’**). **C** Quantification of microtubule fluorescence intensity in placodal cells in control embryos and *UAS-Patronin-RNAi x DaGal4* embryos. Shown are mean +/-SD, note the absence of a significant difference as established by Mann-Whitney test as p=0.3935. **D-E’’’** Patronin-RFP colocalises with a pool of Katanin80-YFP in cells near the invagination pit. The boxed area in **D** is highlighted in **E-E’’’**, arrowheads point to colocalisation. Placode boundaries are indicated by dotted lines and the invagination point by an asterisk.

## Methods

### Drosophila stocks and genetics

The following fly stocks were used in this study: from Bloomington Stock Centre: *Daughterless-Gal4* (*Da-Gal4*; #27608); *UAS-Patronin-RNAi (y1 sc* v1; P{TRiP.HMS01547}attP2)* #36659); *ncd-γ-tubulin-EGFP* (*w1118; P{ncd-γTub37C.GFP}F13F3)*(#56831). Furthermore *Katanin80-YFP (w1118 PBac{602.P.SVS-1}kat80CPTI000764)* [Kyoto Stock Centre; CPTI 000764]; *fkh-Gal4* on chromosome III ^53, 54^; *Asterless-mCherry* (*w; eAsl-mch/Cyo; MKRS/TM6b*) [gift form Jordan Raff]; *YFP-Cnb (w1118; pUbi-YFP-Cnb*) ^29^; *RFP-Cnn* ^30^; *Ubi-EB1-GFP* ^55^; *Patronin-RFP (Patronin-TagRFPattp40[22H02-C])* ^56^; *Patronin-YFP (w1118; Patronin-YFP/Cyo*) ^18^; *GFP-Polo* ^57^; *Sas-4-GFP* ^58^; *Scribble-GFP (w; P{PTT-GA}scribCA07683*) ^59^; *Spd-2-GFP* ^60^; *sqh-TagRFPt[9B]* ^31^; *UAS-deGradFP (w; If/Cyo; UAS>NSlmb-vhhGFP4/TM6b*) ^38^; *UAS-Spastin* on X ^61^.

The following combinations of transgenes were generated in this study:

*sqh-TagRFPt[9B];*; *ncd::γ-tubulin-EGFP*

*Katanin80-YFP;*; *fkh-Gal4*

*Katanin80-YFP;*; *UAS-degradFP*

*Katanin80-YFP; Patronin-RFP; fkh-Gal4*

*Katanin80-YFP; Patronin-RFP; UAS-degradFP*

*Patronin-YFP; Da-Gal4*

*Patronin-YFP; fkh-Gal4*

*UAS-spastin; Patronin-YFP*

*Patronin-YFP; Ubi-TagRFP*

Genotypes analysed are indicated in the figure panels and legends, and are also described under Fly Husbandry below.

### Fly husbandry

To analyse cell boundaries compared to centrosome components, *Asl-mCherry* flies were crossed to *Scribble-GFP*. To analyse microtubule nucleation from Cnn-positive centrosomes, RFP-*Cnn* flies were crossed to *Ubi::EB1-GFP* flies.

To degrade Katanin-YFP specifically in the salivary gland placode, virgins of the genotype *Katanin-YFP fkhGal4* were crossed to males of the genotype *Katanin-YFP UAS-degradFP III*, so that all offspring was homozygous for *Katanin-YFP. Katanin-YFP degradFP III* was analysed as a control.

To analyse Patronin localisation when Katanin-YFP was degraded specifically in the salivary gland placode, virgins of the genotype *Katanin-YFP; Patronin-RFP; fkhGal4* were crossed to males of the genotype *Katanin-YFP; Patronin-RFP; UAS-degradFP*, so that all offspring was homozygous for *Katanin-YFP. Katanin-YFP; Patronin-RFP; degradFP* was analysed as a control.

To analyse Patronin-YFP localisation when microtubules were depleted, virgins of *Patronin-YFP; fkhGal4* were crossed to males of *UAS-Spastin (X); Patronin-YFP*. Using anti-tubulin immunofluorescence, successful MT-depletion was confirmed and only these placodes analysed.

To analyse loss of Patronin in embryos, *UAS-Patronin-RNAi* was expressed throughout the early embryo using *daughterlessGal-4* (*DaGal4*) at 29°C. *UAS-Patronin-RNAi* virgins were crossed to *Patronin-YFP; DaGal4* males. Control embryos were obtained by crossing *Patronin-YFP* males with *DaGal4* virgins. Only embryos where Patronin-YFP staining reduction was visible were analysed.

### Embryo Immunofluorescence Labelling, Confocal, and Live Analysis

Embryos were collected on apple juice-juice plates and processed for immunofluorescence using standard procedures. Briefly, embryos were dechorionated in 50% bleach, fixed in 4% MeOH-free formaldehyde, and devitellinised in a 50% mix of 90% EtOH and Heptane. They were then stained with phalloidin or primary and secondary antibodies in PBT (PBS plus 0.5% bovine serum albumin and 0.3% Triton X-100). anti-E-Cadherin (DCAD2), anti-CrebA (CrebA Rbt-PC) and anti-Crumbs (Cq4) antibodies were obtained from the Developmental Studies Hybridoma Bank at the University of Iowa; anti-aPKC (sc-216; Santa Cruz) anti tyrosinated α-tubulin (YL1/2; ^62^); anti acetylated α-tubulin (clone 6-11B-1; Sigma); anti-α-tubulin (DM1A, Sigma); anti γ-tubulin (clone GTU-88; Sigma); anti-phospho-Histone H3 [Ser10] (Cell Signalling Technology; #9701) anti-Asterless and anti-Cnn ^63^ were a kind gift from Jordan Raff ^64^. Secondary antibodies used were Alexa Fluor 350 (Invitrogen) and Alexa Fluor647 coupled (Jackson ImmunoResearch Laboratories) and Cy3 and Cy5 coupled (Jackson ImmunoResearch Laboratories), and rhodamine-phalloidin was from Thermofisher (R415). Samples were embedded in Vectashield (Vectorlabs H-1000).

Images of fixed samples were acquired on an Olympus FluoView 1200 or a Leica SP8 inverted microscope equipped with 405nm laser line for four-colour imaging as z-stacks to cover the whole apical surface of cells in the placode. Z-stack projections were assembled in ImageJ or Imaris (Bitplane), 3D rendering was performed in Imaris.

For live time-lapse analysis of *Ubi::EB1-GFP RFP-Cnn* embryos and live imaging of *Patronin-YFP; Ubi-TagRFP*, the embryos were dechorionated in 50% bleach, rinsed in water and attached to a coverslip with the ventral side up using heptane glue and covered with Halocarbon Oil 27. Time-lapse sequences of *Ubi::EB1-GFP RFP-Cnn* were acquired on a Leica SP8 inverted microscope (63x/1.4NA Oil objective) as z-stacks, while *Patronin-YFP; Ubi-TagRFP* was imaged on a Zeiss 780 inverted microscope with a 40x/1.3NA Oil objective as a single confocal slice, using linear unmixing to remove the background fluorescence of the embryonic vitelline membrane. Z-stack projections to generate movies were assembled in ImageJ or Imaris.

### Quantifications

#### Proportion of Cnn-positive centrosome nucleating microtubules

Nucleation was manually assessed on time-projections of confocal images from time-lapse movies, assessing whether EB1-GFP comets were seen emanating from Cnn-positive centrosomes or not.

#### Centrosome asymmetry index

Centrosome intensities were measured for both centrosomes in every single cell in an area of 32.48 µm x 25.35 μm close to the invagination pit in late stage 11-early stage 12 embryos on z-projections of the most apical planes (1 to 3 μm depending on the orientation of each embryo). The fluorescence was measured in a circle of 0.68 μm diameter, aiming to cover the maximum area of a centrosome as determined by the area measured for the extension of γ-tubulin, the most outer PCM component of the centrosome ^65^. The ratio was obtained by dividing the fluorescence intensity of the brighter of the two centrosomes by the fluorescence intensity of the other centrosome in any given cell. Numbers analysed are as follow: *Sas4-GFP* (5 placodes, 170 cells); *Spd2-GFP* (4 placodes, 139 cells); α-Asl (9 placodes, 302 cells); α-γ-tubulin (12 placodes, 487 cells); *GFP-Polo* (7 placodes, 270 cells); α-Cnn (9 placodes, 309 cells); *YFP-Cnb* (5 placodes, 157 cells); *ncd::γ-tubulin-EGFP* (8 placodes, 284 cells).

#### MT intensity measurements

The raw intensity of microtubule labeling in γ-tubulin-EGFP overexpression or Patronin-YFP depletion experiments (α-tubulin, anti-tyrosinated α-tubulin or anti-acetylated α-tubulin) was measured in in the entire placode on z-projections of the most apical planes (1 to 3 μm depending on the orientation of each embryo).

#### Automated cell segmentation and apical area analysis

For the analysis of apical cell area, images of fixed embryos of late stage 11/early stage 12 placodes, labelled with DE-Cadherin to highlight cell membranes and with dCrebA to mark salivary gland fate, were analysed. Cells were segmented in confocal image stacks, with cell analysis software as used previously ^8, 66-68^. Briefly, the shape of the curved placode surface was identified in each z-stack as a contiguous ‘blanket’ spread over the cortical signal. Quasi-2D images for cell segmentation containing clear cell cortices were extracted as a maximum intensity projection of the 1 or 1.5 µm thick layer of tissue below the blanket. These images were segmented using an adaptive watershed algorithm. Manual correction was used to perfect cell outlines for fixed embryos. Only cells of the salivary placode were used in subsequent analyses and were distinguished based on dCrebA staining.

#### Number of wavy junctions

The fraction of junctions displaying waviness was determined manually by counting how many junctions within the secretory region of the analysed placodes (n=7 placodes for control, n=10 placodes for *ncd::γ-tubulin-EGFP* embryos) out of the total number of junctions in that area displayed waviness, i.e. an undulating deviation from the direct line between two vertices.

#### Straightness measurements of junctions

This straightness of a junction was determined by dividing the ideal junction length between vertices (L_[direct route]_) by the actual junction path length (L_[junction]_, see Fig. 3K), based on the E-cadherin staining as in ^32^. L_[direct route]_ and L_[junction]_ were determined manually using FIJI. Only junctions considered as wavy junctions in the above quantification were quantified (n=90) in *ncd::γ-tubulin-EGFP* embryos that overexpress γ-tubulin-EGFP. The same number of junctions was analysed in control placodes. As wavy junctions were also at times observed in control embryos and in order to achieve unbiased analysis, junctions to be quantified in control embryos were selected using an online random number generator.

#### Katanin80-YFP accumulation at centrosomes

The Katanin80-YFP protein trap strain displays a strong homogeneous background fluorescence. Due to this, images were processed as follows to be able to determine actual Katanin80-YFP fluorescence at centrosomes rather than measure the background fluorescence. The Katanin80-YFP channel was selected and subjected to background subtraction using the subtract background function in Fiji with a rolling ball radius of 50 pixels. The image was then smoothed and thresholded using the automatic Triangle method in Fiji to obtain a mask of the Katanin80-YFP channel. The pixel values of the mask were divided by 255 to generate a binary mask of Katanin80-YFP where Katanin signal corresponds to pixel value of 1. The binary mask was then used to multiply the pixel values of the original image by 0 or 1, to obtain the actual Katanin80-YFP staining corresponding to pixels with a value of 1 in the binary mask, and to therefore obtain the processed Katanin80-YFP image. Katanin80-YFP intensity was then measured in the processed image in an area of 33 x 21μm close to the invagination pit, by drawing a circle ROI of 0.68 μm diameter surrounding each centrosome marked by Asl-mCherry. Results shown correspond to 232 cells in 7 embryos.

#### Quantification of Katanin80-YFP

Katanin80-YFP staining was measured in the placode and the surrounding epidermis (33 control embryos and 34 Katanin-depleted embryos were analysed in z-projections of the most apical planes, 1 to 3 μm depending on the orientation of each embryo). As Katanin80-YFP embryos displayed a hazy background fluorescence, we determined the background fluorescence from five ROIs deeper within the embryo where there is no *bona fide* Katanin fluorescence and subtracted this from the values obtained. We then calculated the ratio of Katanin80-YFP enrichment in the placode by dividing the intensity value in the placode by the intensity value determined in the surrounding epidermis. 33 control embryos and 34 Katanin-depleted embryos were analysed.

#### Acetylated α-tubulin accumulation near centrosomes

The microtubule intensity was measured in an ROI of 0.68 μm diameter around the centrosomes (labelled with Asl-mCherry) on Z projections of the most apical planes (1 to 3 μm depending on the orientation of each embryo), and the intensity was normalised against the mean microtubule intensity in the entire placode.

#### Medial accumulation of phalloidin, sqh-RFP, Patronin-YFP and Patronin-RFP

Images were taken of salivary gland placodes and surrounding tissue at late stage 11/early stage 12. Maximum intensity projections of the apical surface of placodal cells were generated using 3-5 optical section separated by 1 µm each in z. For each embryo analysed, fluorescence measurements were made for all secretory cells within the placode except cells close to the actomyosin cable. The medial and junctional values were measured after drawing the cells outlines (7-pixel wide line) with a home-made plugin in Fiji, available on request. 10 cells were similarly analysed in the surrounding tissue. The graphs depict the enrichment of medial intensity compared to the surrounding epidermis obtained by dividing individual values measured in the placode by the mean intensity of the cells in the surrounding epidermis.

For Patronin-YFP, Sqh-RFP and phalloidin quantifications, the graphs display the medial accumulation corresponding to the ratio between the medial intensity for each cell in the placode divided by the mean intensity of the 10 cells outside the placode. For the comparison of Patronin-YFP in the placode and the surrounding epidermis, the graph displays the ratio between medial Patronin-YFP intensity versus the background intensity measured deeper within the embryo where there is no *bona fide* Patronin fluorescence. For Patronin-RFP, due to the noisy labeling in the surrounding epidermis, the graph displays the ratio between Patronin-RFP medial staining versus junctional staining for each cell in the placode.

### Statistical Analysis

Significance was determined using two-tailed Student’s *t*-test, non-parametric Mann-Whitney test for non-Gaussian distribution, unpaired t-test with Welch’s correction for data with unequal standard deviations, and Kolmogorov-Smirnov (K-S) test for the comparison of cumulative distributions. For the γ-tubulin overexpression experiment (Fig.3), the K-S test did not show any significant difference for the cumulative data between the control and γ-tubulin-overexpressing embryos. However, comparisons of cell apical areas in small size windows revealed a significant difference in the distribution of cells with the smallest apical areas (0 μm^2^ < apical area < 5 μm^2^, p=0.0361; 5 μm^2^ < apical area < 10 μm^2^, p=0.0361). Centrosome asymmetries (Fig. 2) were compared using a Kruskal-Wallis test (non-parametric one-way ANOVA) for multiple pairwise comparisons. Results were considered significant when P<0.05. N values and statistical tests used are indicated in the Results section as well as the Figure Legends.

## References

1. Petry, S. & Vale, R.D. Microtubule nucleation at the centrosome and beyond. Nat Cell Biol 17, 1089–1093 (2015).

2. Kollman, J.M., Merdes, A., Mourey, L. & Agard, D.A. Microtubule nucleation by gamma-tubulin complexes. Nat Rev Mol Cell Biol 12, 709–721 (2011).

3. Tas, R.P. et al. Differentiation between Oppositely Oriented Microtubules Controls Polarized Neuronal Transport. Neuron 96, 1264–1271 e1265 (2017).

4. Zenker, J. et al. A microtubule-organizing center directing intracellular transport in the early mouse embryo. Science 357, 925–928 (2017).

5. Martin, M., Veloso, A., Wu, J., Katrukha, E.A. & Akhmanova, A. Control of endothelial cell polarity and sprouting angiogenesis by non-centrosomal microtubules. eLife 7 (2018).

6. Ko, C.S., Tserunyan, V. & Martin, A.C. Microtubules promote intercellular contractile force transmission during tissue folding. J Cell Biol 218, 2726–2742 (2019).

7. Singh, A. et al. Polarized microtubule dynamics directs cell mechanics and coordinates forces during epithelial morphogenesis. Nat Cell Biol 20, 1126–1133 (2018).

8. Booth, A.J., Blanchard, G.B., Adams, R.J. & Röper, K. A dynamic microtubule cytoskeleton directs medial actomyosin function during tube formation. Dev Cell 29, 562–576 (2014).

9. Toya, M. et al. CAMSAP3 orients the apical-to-basal polarity of microtubule arrays in epithelial cells. Proc Natl Acad Sci U S A 113, 332–337 (2016).

10. Pope, K.L. & Harris, T.J. Control of cell flattening and junctional remodeling during squamous epithelial morphogenesis in Drosophila. Development 135, 2227–2238 (2008).

11. Garcia De Las Bayonas, A., Philippe, J.M., Lellouch, A.C. & Lecuit, T. Distinct RhoGEFs Activate Apical and Junctional Contractility under Control of G Proteins during Epithelial Morphogenesis. Curr Biol 29, 3370–3385 e3377 (2019).

12. Feldman, J.L. & Priess, J.R. A role for the centrosome and PAR-3 in the hand-off of MTOC function during epithelial polarization. Curr Biol 22, 575–582 (2012).

13. Sanchez-Corrales, Y.E., Blanchard, G.B. & Roper, K. Radially patterned cell behaviours during tube budding from an epithelium. eLife 7 (2018).

14. Foe, V.E. & Alberts, B.M. Studies of nuclear and cytoplasmic behaviour during the five mitotic cycles that precede gastrulation in Drosophila embryogenesis. J Cell Sci 61, 31–70 (1983).

15. Edgar, B.A. & O’Farrell, P.H. Genetic control of cell division patterns in the Drosophila embryo. Cell 57, 177–187 (1989).

16. Callaini, G., Whitfield, W.G. & Riparbelli, M.G. Centriole and centrosome dynamics during the embryonic cell cycles that follow the formation of the cellular blastoderm in Drosophila. Exp Cell Res 234, 183–190 (1997).

17. Hendershott, M.C. & Vale, R.D. Regulation of microtubule minus-end dynamics by CAMSAPs and Patronin. Proc Natl Acad Sci U S A 111, 5860–5865 (2014).

18. Nashchekin, D., Fernandes, A.R. & St Johnston, D. Patronin/Shot Cortical Foci Assemble the Noncentrosomal Microtubule Array that Specifies the Drosophila Anterior-Posterior Axis. Dev Cell 38, 61–72 (2016).

19. Stevens, N.R., Raposo, A.A., Basto, R., St Johnston, D. & Raff, J.W. From stem cell to embryo without centrioles. Curr Biol 17, 1498–1503 (2007).

20. Basto, R. et al. Flies without centrioles. Cell 125, 1375–1386 (2006).

21. Chandrasekaran, V. & Beckendorf, S.K. Tec29 controls actin remodeling and endoreplication during invagination of the Drosophila embryonic salivary glands. Development 132, 3515–3524 (2005).

22. Dobbelaere, J. et al. A genome-wide RNAi screen to dissect centriole duplication and centrosome maturation in Drosophila. PLoS biology 6, e224 (2008).

23. Lerit, D.A. et al. Interphase centrosome organization by the PLP-Cnn scaffold is required for centrosome function. J Cell Biol 210, 79–97 (2015).

24. Conduit, P.T. et al. A molecular mechanism of mitotic centrosome assembly in Drosophila. eLife 3, e03399 (2014).

25. Fu, J., Hagan, I.M. & Glover, D.M. The centrosome and its duplication cycle. Cold Spring Harb Perspect Biol 7, a015800 (2015).

26. Zhang, J. & Megraw, T.L. Proper recruitment of gamma-tubulin and D-TACC/Msps to embryonic Drosophila centrosomes requires Centrosomin Motif 1. Mol Biol Cell 18, 4037–4049 (2007).

27. Ramani, A. et al. Plk1/Polo Phosphorylates Sas-4 at the Onset of Mitosis for an Efficient Recruitment of Pericentriolar Material to Centrosomes. Cell reports 25, 3618–3630 e3616 (2018).

28. Januschke, J. et al. Centrobin controls mother-daughter centriole asymmetry in Drosophila neuroblasts. Nat Cell Biol 15, 241–248 (2013).

29. Januschke, J., Llamazares, S., Reina, J. & Gonzalez, C. Drosophila neuroblasts retain the daughter centrosome. Nature communications 2, 243 (2011).

30. Conduit, P.T. et al. Centrioles regulate centrosome size by controlling the rate of Cnn incorporation into the PCM. Curr Biol 20, 2178–2186 (2010).

31. Ambrosini, A., Rayer, M., Monier, B. & Suzanne, M. Mechanical Function of the Nucleus in Force Generation during Epithelial Morphogenesis. Dev Cell 50, 197–211 e195 (2019).

32. Sumi, A. et al. Adherens Junction Length during Tissue Contraction Is Controlled by the Mechanosensitive Activity of Actomyosin and Junctional Recycling. Dev Cell 47, 453–463 e453 (2018).

33. Balaji, R., Weichselberger, V. & Classen, A.K. Response of Drosophila epithelial cell and tissue shape to external forces in vivo. Development 146 (2019).

34. Brodu, V., Baffet, A.D., Le Droguen, P.M., Casanova, J. & Guichet, A. A developmentally regulated two-step process generates a noncentrosomal microtubule network in Drosophila tracheal cells. Dev Cell 18, 790–801 (2010).

35. Sanchez, A.D. & Feldman, J.L. Microtubule-organizing centers: from the centrosome to non-centrosomal sites. Curr Opin Cell Biol 44, 93–101 (2017).

36. Muroyama, A. & Lechler, T. Microtubule organization, dynamics and functions in differentiated cells. Development 144, 3012–3021 (2017).

37. McNally, F.J. & Vale, R.D. Identification of katanin, an ATPase that severs and disassembles stable microtubules. Cell 75, 419–429 (1993).

38. Caussinus, E., Kanca, O. & Affolter, M. Fluorescent fusion protein knockout mediated by anti-GFP nanobody. Nature structural & molecular biology 19, 117–121 (2012).

39. Yvon, A.M. & Wadsworth, P. Non-centrosomal microtubule formation and measurement of minus end microtubule dynamics in A498 cells. J Cell Sci 110 (Pt 19), 2391–2401 (1997).

40. Akhmanova, A. & Hoogenraad, C.C. Microtubule minus-end-targeting proteins. Curr Biol 25, R162–171 (2015).

41. Goodwin, S.S. & Vale, R.D. Patronin regulates the microtubule network by protecting microtubule minus ends. Cell 143, 263–274 (2010).

42. Jiang, K. et al. Microtubule minus-end stabilization by polymerization-driven CAMSAP deposition. Dev Cell 28, 295–309 (2014).

43. Meng, W., Mushika, Y., Ichii, T. & Takeichi, M. Anchorage of microtubule minus ends to adherens junctions regulates epithelial cell-cell contacts. Cell 135, 948–959 (2008).

44. Chen, J.V., Buchwalter, R.A., Kao, L.R. & Megraw, T.L. A Splice Variant of Centrosomin Converts Mitochondria to Microtubule-Organizing Centers. Curr Biol 27, 1928–1940 e1926 (2017).

45. Ahmad, F.J., Yu, W., McNally, F.J. & Baas, P.W. An essential role for katanin in severing microtubules in the neuron. J Cell Biol 145, 305–315 (1999).

46. Le, T.P. & Chung, S.Y. Microtubule-dependent protein trafficking promotes apical constriction during tissue invagination. bioRxiv https://www.biorxiv.org/content/10.1101/827378v1 (2019).

47. Ingber, D.E. Tensegrity I. Cell structure and hierarchical systems biology. J Cell Sci 116, 1157–1173 (2003).

48. Dong, C. et al. CAMSAP3 accumulates in the pericentrosomal area and accompanies microtubule release from the centrosome via katanin. J Cell Sci 130, 1709–1715 (2017).

49. Khanal, I., Elbediwy, A., Diaz de la Loza Mdel, C., Fletcher, G.C. & Thompson, B.J. Shot and Patronin polarise microtubules to direct membrane traffic and biogenesis of microvilli in epithelia. J Cell Sci 129, 2651–2659 (2016).

50. Noordstra, I. et al. Control of apico-basal epithelial polarity by the microtubule minus-end-binding protein CAMSAP3 and spectraplakin ACF7. J Cell Sci 129, 4278–4288 (2016).

51. Röper, K. Anisotropy of Crumbs and aPKC Drives Myosin Cable Assembly during Tube Formation. Dev Cell 23, 939–953 (2012).

52. Brand, A.H. & Perrimon, N. Targeted gene expression as a means of altering cell fates and generating dominant phenotypes. Development 118, 401–415 (1993).

53. Henderson, K.D. & Andrew, D.J. Regulation and function of Scr, exd, and hth in the Drosophila salivary gland. Dev Biol 217, 362–374 (2000).

54. Zhou, B., Bagri, A. & Beckendorf, S.K. Salivary gland determination in Drosophila: a salivary-specific, fork head enhancer integrates spatial pattern and allows fork head autoregulation. Dev Biol 237, 54–67 (2001).

55. Gallaud, E. et al. Ensconsin/Map7 promotes microtubule growth and centrosome separation in Drosophila neural stem cells. J Cell Biol 204, 1111–1121 (2014).

56. Takeda, M., Sami, M.M. & Wang, Y.C. A homeostatic apical microtubule network shortens cells for epithelial folding via a basal polarity shift. Nat Cell Biol 20, 36–45 (2018).

57. Moutinho-Santos, T., Sampaio, P., Amorim, I., Costa, M. & Sunkel, C.E. In vivo localisation of the mitotic POLO kinase shows a highly dynamic association with the mitotic apparatus during early embryogenesis in Drosophila. Biol Cell 91, 585–596 (1999).

58. Novak, Z.A., Conduit, P.T., Wainman, A. & Raff, J.W. Asterless licenses daughter centrioles to duplicate for the first time in Drosophila embryos. Curr Biol 24, 1276–1282 (2014).

59. Buszczak, M. et al. The carnegie protein trap library: a versatile tool for Drosophila developmental studies. Genetics 175, 1505–1531 (2007).

60. Dix, C.I. & Raff, J.W. Drosophila Spd-2 recruits PCM to the sperm centriole, but is dispensable for centriole duplication. Curr Biol 17, 1759–1764 (2007).

61. Sherwood, N.T., Sun, Q., Xue, M., Zhang, B. & Zinn, K. Drosophila spastin regulates synaptic microtubule networks and is required for normal motor function. PLoS biology 2, e429 (2004).

62. Kilmartin, J.V., Wright, B. & Milstein, C. Rat monoclonal antitubulin antibodies derived by using a new nonsecreting rat cell line. J Cell Biol 93, 576–582 (1982).

63. Lucas, E.P. & Raff, J.W. Maintaining the proper connection between the centrioles and the pericentriolar matrix requires Drosophila centrosomin. J Cell Biol 178, 725–732 (2007).

64. Stevens, N.R., Dobbelaere, J., Wainman, A., Gergely, F. & Raff, J.W. Ana3 is a conserved protein required for the structural integrity of centrioles and basal bodies. J Cell Biol 187, 355–363 (2009).

65. Fu, J. & Glover, D.M. Structured illumination of the interface between centriole and peri-centriolar material. Open Biol 2, 120104 (2012).

66. Blanchard, G.B. et al. Tissue tectonics: morphogenetic strain rates, cell shape change and intercalation. Nat Methods 6, 458–464 (2009).

67. Blanchard, G.B., Murugesu, S., Adams, R.J., Martinez-Arias, A. & Gorfinkiel, N. Cytoskeletal dynamics and supracellular organisation of cell shape fluctuations during dorsal closure. Development 137, 2743–2752 (2010).

68. Butler, L.C. et al. Cell shape changes indicate a role for extrinsic tensile forces in Drosophila germ-band extension. Nat Cell Biol 11, 859–864 (2009).

